# The cytokinesis associated proteins CITK and ASPM-1 regulate neuronal microtubule dynamics and polarity in *C. elegans*

**DOI:** 10.1101/2025.04.24.650367

**Authors:** Sunanda Sharma, Keerthana Ponniah, Anindya Ghosh-Roy

**Affiliations:** Department of Cellular & Molecular Neuroscience, National Brain Research Centre, Manesar, Haryana-122052, India; Current Affiliation: Clem Jones Centre for Ageing Dementia Research, Queensland Brain Institute, The University of Queensland, Brisbane, QLD 4072, Australia

**Keywords:** *C. elegans*, microtubule dynamics, microtubule polarity, touch neuron, *klp-7*, *citk-1*, *citk-2*, *aspm-1*

## Abstract

The polarized architecture of neurons is intricately associated with modulation of microtubule dynamics. Over the years several microtubule-associated-factors that modulate neuronal polarity have been identified. However, the precise details of how microtubule arrangement and stability is established in axon and dendrites is not clearly understood. To uncover relevant factors involved in the biological pathways governing microtubule regulation in neuron, we conducted a suppressor screen using the neuronal ectopic extension phenotype caused due to loss of kinesin-13 family microtubule depolymerizing protein KLP-7 in *C. elegans*. Interestingly, apart from eleven variants of α (*mec-12*) and β (*mec-7*) tubulins, we isolated a variant of cytokinesis associated protein, *W02B8.2/citk-1,* the kinase-less worm orthologue of mammalian citron-rho interacting kinase (CIT). Little is known about the role of CITK in microtubule regulation in post-mitotic neurons. In this study, we found that the kinase-less worm orthologues of CIT, *citk-1* and *citk-2* redundantly modulate microtubule stability in the axon-like anterior process and maintain the population of plus-end-out microtubules in the dendrite-like posterior process of the PLM mechanosensory neurons in a cell autonomous manner. In the absence of *citk-1*, PLM neurons exhibit variable morphological defects including defects in migration, growth, and guidance. Moreover, the neuronal and microtubule phenotypes of loss of both *citk-1* and *citk-2* were phenocopied by the mutant animals of *aspm-1,* the worm homolog of abnormal spindle-like microcephaly-associated protein (ASPM), suggesting a genetic association, similar to their association in dividing mammalian cells. These observations suggest that the cytokinesis associated citron kinase and ASPM-1 have non-mitotic roles in *C. elegans* mechanosensory neurons in the regulation of microtubules.

## Introduction

The fundamental unit of nervous system, a neuron, is a highly polarized cell. Its ability to transmit information unidirectionally over long distances in neural networks relies on its structural and functional compartmentalization into dendrites and axons, the information receiving and transmitting processes (1). One of the most sought out questions in neurobiology is how this elaborate architecture is developed and maintained throughout the lifespan of an organism. Previous studies have demonstrated that the regulation of neuronal polarity is closely linked to the arrangement of microtubule polarity and dynamics within neuronal compartments (2–6). Microtubules are tubular, hollow, heteropolymers of α-β tubulin dimers, arranged in a head-to-tail manner rendering them an intrinsic polarity (7, 8). The visibly dynamic end with a β-tubulin exposed is called plus end and the less dynamic end with an α-tubulin exposed is called minus-end (9). In the neuronal system, axons have a unipolar plus-end-out (plus-end-distal to cell body) microtubule arrangement, while dendrites have a mixed microtubule polarity in vertebrates and a minus-end-out polarity in the invertebrate neurons (10, 11). However, little is known about how this compartmentalized microtubule arrangement and dynamics is achieved in neurons.

Over the years several microtubule associated factors including microtubule nucleation factors (such as y-TURC), polymerization factors (for instance, tubulins, CRMP-2/UNC-33), microtubule anchoring proteins (TRIM46, UNC-44/Ankyrin-G) and regulators of microtubule-end dynamics (plus-end-binding proteins EBP-2/3, minus-end binding protein Patronin/CAMSAP2/3) have been identified as regulators of neuronal microtubule dynamics and arrangement (6, 12–16). The microtubule cross-linking and anchoring factors such as TRIM46 and Ankyrin-G organize the microtubules in parallel bundles with plus-end-out polarity in the axon initial segment (17–20). The microtubule end binding proteins act as microtubule stabilizing factors in axon and dendrites (21–23). While Kinesin-13 family microtubule depolymerizing factor KIF2A/KLP-7 has been shown to be a gate keeper of dendritic identity in mouse, fish and *C. elegans* neurons (2, 4, 5, 24–26). In the absence of kinesin-13, microtubules are hyper-stabilized leading to reorganization of dendritic microtubules in a plus end out orientation (2, 25, 26). However, it’s not completely clear how the microtubules in the distal axon, which is farther away from the axon initial segment, remain bundled and optimally stabilized. Furthermore, our understanding of how microtubules retain a mixed polarity organization within dendritic compartments is fragmentary.

The mechanosensory neurons of *C. elegans* is an excellent model to study the regulation of microtubule dynamics and polarity in neurons (27–29). Several microtubule regulating factors as well as molecules regulating post-translational modifications of tubulins have been identified and studied in the ALM and PLM mechanosensory neurons (20, 25, 30–36). The PLM (posterior lateral microtubule) neuron is a well characterized bipolar neuron that under the guidance of Wnt signaling gradient polarizes and extends its neurites in the antero-posterior axis (2, 37–40). Its anterior neurite exhibits plus-end out microtubule arrangement, similar to vertebrate axon, while the posterior short neurite displays a mixed microtubule polarity, similar to vertebrate dendrites. Any perturbation in the microtubule dynamics may lead to distinct morphological changes in these neurons. For instance, loss of Kinesin-13 family microtubule depolymerizing factor KLP-7 results in hyper-stabilization of microtubules and hence, ectopic neurites sprouting from mechanosensory neurons of mutant animals including overgrowth of PLM posterior process. The ALM (anterior lateral microtubule) neurons on the other hand, are largely unipolar, however, in *klp-7*(*0*) animals ALM neurons develop a visibly striking posterior ectopic extension (25). These ectopic growths can be suppressed by destabilizing microtubules (2, 30). We hypothesized that the *klp-7*(*0*) mutant phenotype, combined with the powerful genetic toolbox of *C. elegans* (41), could be a promising platform for designing a suppressor screen aimed at identifying key regulators of neuronal microtubule dynamics which could enhance our understanding of how microtubule dynamics and polarity are regulated in neurons.

In this study, we successfully isolated twenty-six suppressor variants in a screen and mapped twelve of these to the loci of mechanosensory neuron specific tubulins *mec-12* (α-tubulin) and *mec-7* (β-tubulin). Incidentally, we isolated a variant of *W02B8.2*/*citk-1,* a kinase-less worm orthologue of mammalian citron-rho kinase-associated protein/CIT/MCPH17 (autosomal recessive primary microcephaly-17). Live imaging of EBP-2::GFP reporter revealed that *citk-1,* along with its paralog *citk-2* reigns in the dynamic microtubules in the axon-like anterior process of PLM mechanosensory neurons and maintains the plus-end-out microtubule population in the dendrite-like posterior process in a cell-autonomous manner. Furthermore, loss of both citron kinase proteins results in guidance and morphological defects in these neurons. Additionally, we found that the loss of ASPM-1, worm orthologue of mammalian ASPM (Abnormal Spindle-like Microcephaly-Associated Protein, MCPH5), a known partner of CIT in microtubule regulation during cell division phenocopied the citron kinase mutant animals, suggesting they might be working in the same pathway to regulate microtubule dynamics in the PLM mechanosensory neurons. The findings suggest that cytokinesis associated proteins *citk-1/citk-2* and *aspm-1* have a post-differentiation role in regulation of microtubule dynamics in neurons and hence, neuron morphology and maintenance.

## Results

### A modifier screen using the ectopic extension phenotype of *klp-7*(*0*) identifies neuronal microtubule regulators including tubulin variants

Previous studies have shown that loss of KLP-7, a kinesin-13 family microtubule depolymerizing protein, leads to ectopic neurite extensions in mechanosensory neurons, (2, 25), in particular the strikingly elongated extension of the posterior process of the ALM mechanosensory neurons of *C. elegans* (Figure 1A-B’). The wild type *C. elegans* have two unipolar, bilateral ALM neurons situated a little anterior to the midbody region and characterized by a long anterior neurite that extends anterolaterally parallel to the dorsal nerve cord (Figure 1A-B’). Occasionally, it may extend a short posterior process, shorter than twice the length of the ALM cell body (27, 28). However, in *klp-7(0)* mutant animals this posterior process overgrows and may even extend beyond the cell body of posteriorly situated PVM mechanosensory neuron (Figure 1A-B’). Furthermore, it was previously shown that this phenotype can be suppressed by destabilization of microtubules, either by colchicine treatment or by a second site mutation in mechanosensory neuron specific tubulins, β-tubulin *mec-7*, or α-tubulin *mec-12* (2, 30, 42).

**Figure 1.**
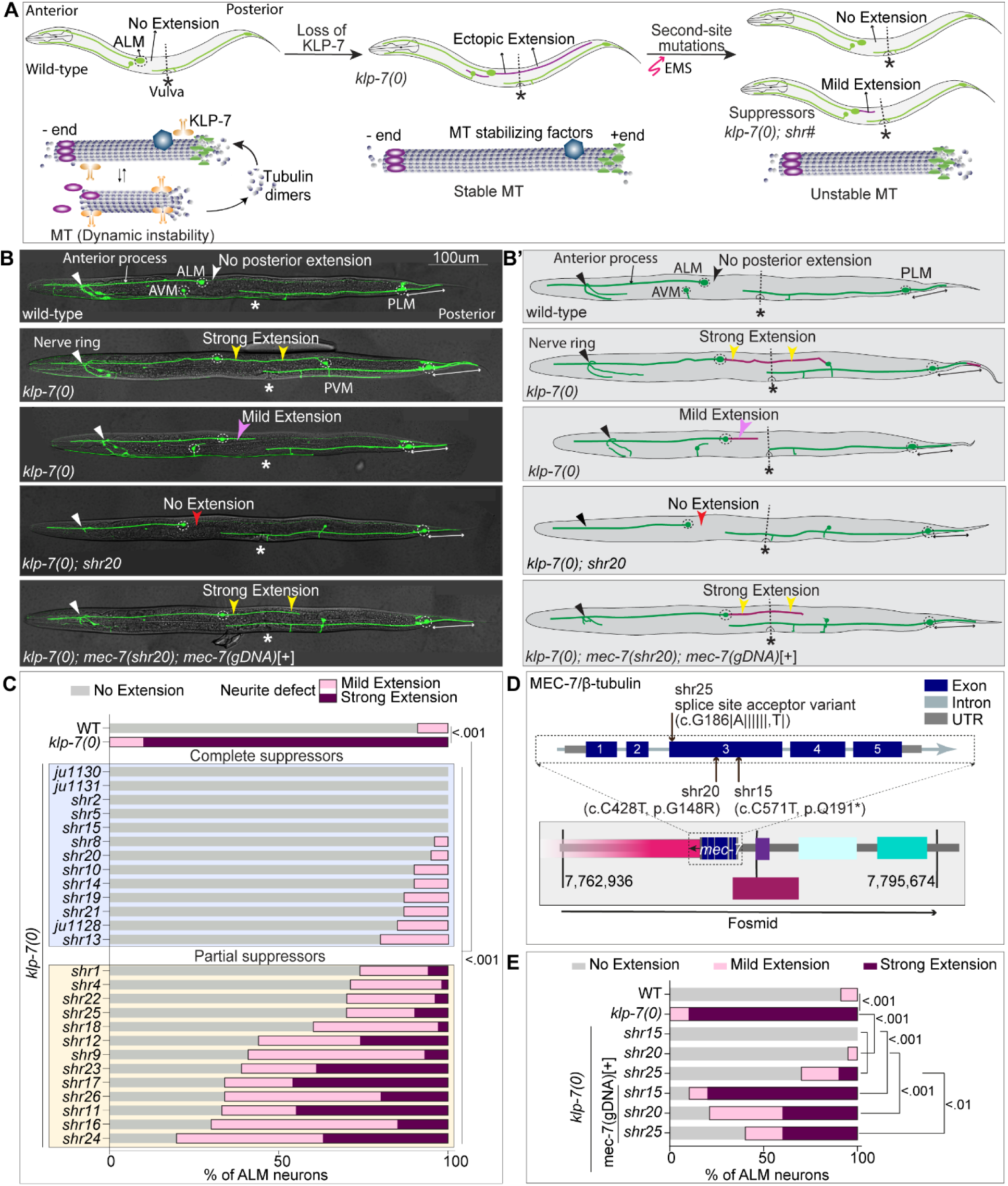
Isolation and mapping of suppressors of *klp-7(0)* ALM ectopic extension phenotype. **(A)** Schematics illustrates the hypothesis of EMS genetic screening. The wild-type animals with a steady-state microtubule dynamics exhibit no posterior extension from ALM mechanosensory neuron. Loss of function mutations in the microtubule depolymerizing protein KLP-7 (orange structure) leads to a posterior ectopic extension (drawn in purple) from the ALM cell body, associated with stable microtubules in these animals. The stable microtubules in the *klp-7(0)* mutant can be destabilized by EMS-induced second-site mutations in other microtubule regulators. The *klp-7(0)* ectopic extension phenotype would be reduced or absent in these suppressor mutant backgrounds. The blue and gray spheres represent α and β tubulins, respectively, which form the structure of microtubules. The purple ovals represent minus-end binding protein and the green hourglass like-structures represents plus-end binding proteins. The deep sky-blue hexagon depicts a probable microtubule stabilizing protein being mutated in suppressor mutant. Microtubule dynamics scheme is adapted from (10). **(B-B’)** Representative confocal images and schematics of ALM and PLM neurons in the L4 staged wild-type, *klp-7(0)* mutant, a suppressor *klp-7(0); mec-7(shr20)* and a *klp-7(0); mec-7(shr20); mec-7* (genomic DNA)[+] animals. The anterior end of animal is placed left, and vulva is positioned downwards in this and following figures. The ALM and PLM mechanosensory neurons of these animals express GFP (green fluorescent protein) under mechanosensory neuron specific promoter, *Pmec-7* (*Pmec-7::GFP, muIs32)*. (0) represents the loss of function deletion allele *(tm2143)* of *klp-7.* The white arrowhead labels the no ectopic extension phenotype in wild-type animal. The yellow and pink arrowheads point to strong and mild posterior extensions from ALM in *klp-7(0)* animals that ends either before or after the vulval position landmark(*), respectively. The red arrowhead points to suppression of ALM ectopic extension phenotype in *klp-7(0); mec-7(shr20)* mutant which is rescued (yellow arrowheads) in *klp-7(0); mec-7(shr20); shrEx396(mec-7(gDNA)*[+]) animals. A double-headed white arrow is drawn along the length of the PLM posterior process. **(C)** Quantification of the percentage of ALM neurons with ectopic extension in the *klp-7(0)* and the suppressor backgrounds *klp-7(0); shr#*, where *shr#* is the allele number. N = 3–4 independent replicates, n (number of neurons) = 45-55. (D) A schematic representation of the fosmid WRM062bA08, which includes the complete genomic cassette of the *mec-7 gene*, as shown. (E) Quantification of the percentage of ALM neurons with ectopic extension in the wild-type, *klp-7(0)*, *klp-7(0); mec-7(shr15*), *klp-7(0); mec-7(shr20), klp-7(0); mec-7(shr25), klp-7(0); mec-7(shr15)*; *mec-7*(gDNA) [+], *klp-7(0); mec*-*7(shr20); mec-7*(gDNA) [+] and *klp-7(0); mec-7(shr25); mec-7*(gDNA) [+] backgrounds, where g stands for genomic and *mec-7*(gDNA) is the fosmid WRM062bA08. N = 3–5 independent replicates, n (number of neurons) = 45-50. For C and E, P values from 2×2 Fisher’s exact test comparing number of animals with No extension phenotype and number of animals with a Neurite defect (mild-extension + strong extension). across different genotypes.

This suggested that by using a modifier screen in this sensitized background of *klp-7(0)* we can isolate variants of novel microtubule regulators as suppressors of ectopic extension phenotype, which can further enhance our understanding of neuronal microtubule regulation (Figure 1A). Hence, we conducted a clonal EMS mutagenesis screening and isolated twenty-six unique suppressor variants from 12,422 F1’s (Figure 1A-B’ and S1A). The progeny of these suppressor mutants was qualitatively scored to determine the extent of suppression. The ectopic extension phenotype for the purpose of this study was classified as strong extension if it reached or crossed the vulval position and mild extension if it ended before vulva, while the wild-type like phenotype was classified as a no extension phenotype (Figure 1B-B’). On the basis of this classification the suppressor variant was categorized as complete suppressor if none of the ALM processes in population showed the strong extension phenotype, including suppressors *shr10, shr15, shr19, shr20*, *shr21* else it was categorized as a mild suppressor, including suppressors *shr1, shr9, shr17, shr22* and *shr25* (Figure 1C).

The next obvious step was to identify the molecular nature of these isolated variants. Using EMS density deep sequencing mapping strategy we mapped three of the isolated suppressor variants to mechanosensory neuron specific β-tubulin, *mec-7*, (*shr15, shr20* and *shr25)* (Figure 1D and S1A-C), and eight to α-tubulin, *mec-12,* (*shr1, shr9, shr10, shr14, shr17, shr19, shr21* and *shr22)* (Figure S1 E-F). The mapping results were further validated by replenishing the mutated protein in the suppressor variant by extra-chromosomal expression of a wild-type copy of the complete genomic casette of candidate gene (*mec-7*, in this case) which could rescue the ectopic extension phenotype of *klp-7(0)* in these suppressor mutant backgrounds (Figure 1D-E).

In summary, we isolated twenty-six unique suppressors of *klp-7(0)* ectopic extension phenotype with varying extent of suppression. Eleven suppressor mutants mapped to mechanosensory neuron specific tubulins *mec-7* and *mec-12.* As discussed previously, it was already known that loss of *mec-7* and *mec-12* suppresses ectopic extension phenotype by destabilizing microtubules (2, 30, 42). Thus, isolation of *mec-7* and *mec-12* variants in this screen was exciting as it suggested that this screen is a promising approach to identify neuronal microtubule regulators to further our understanding of the regulation of microtubules in neurons. And interestingly we did identify some unexpected candidates including an RNA binding protein, *mbl-1* (42) and a cytokinesis associated factor, *citk-1*, which we will be discussing in detail in the following sections.

### Mutations in a previously uncharacterized protein *W02B8.2/citk-1* suppress the ectopic extension phenotype of *klp-7(0)* mutant

We mapped one of the partial suppressors, *shr11,* to a proline to leucine substitution at the 855^th^ amino acid in the genomic locus of a previously uncharacterized protein W02B8.2 on Chromosome II (Figure 2A-B and S2A). A previous report suggested that *W02B8.2* is a homolog of mammalian citron-rho interacting kinase which is required for the organization of midbody during cytokinesis (43).

**Figure 2.**
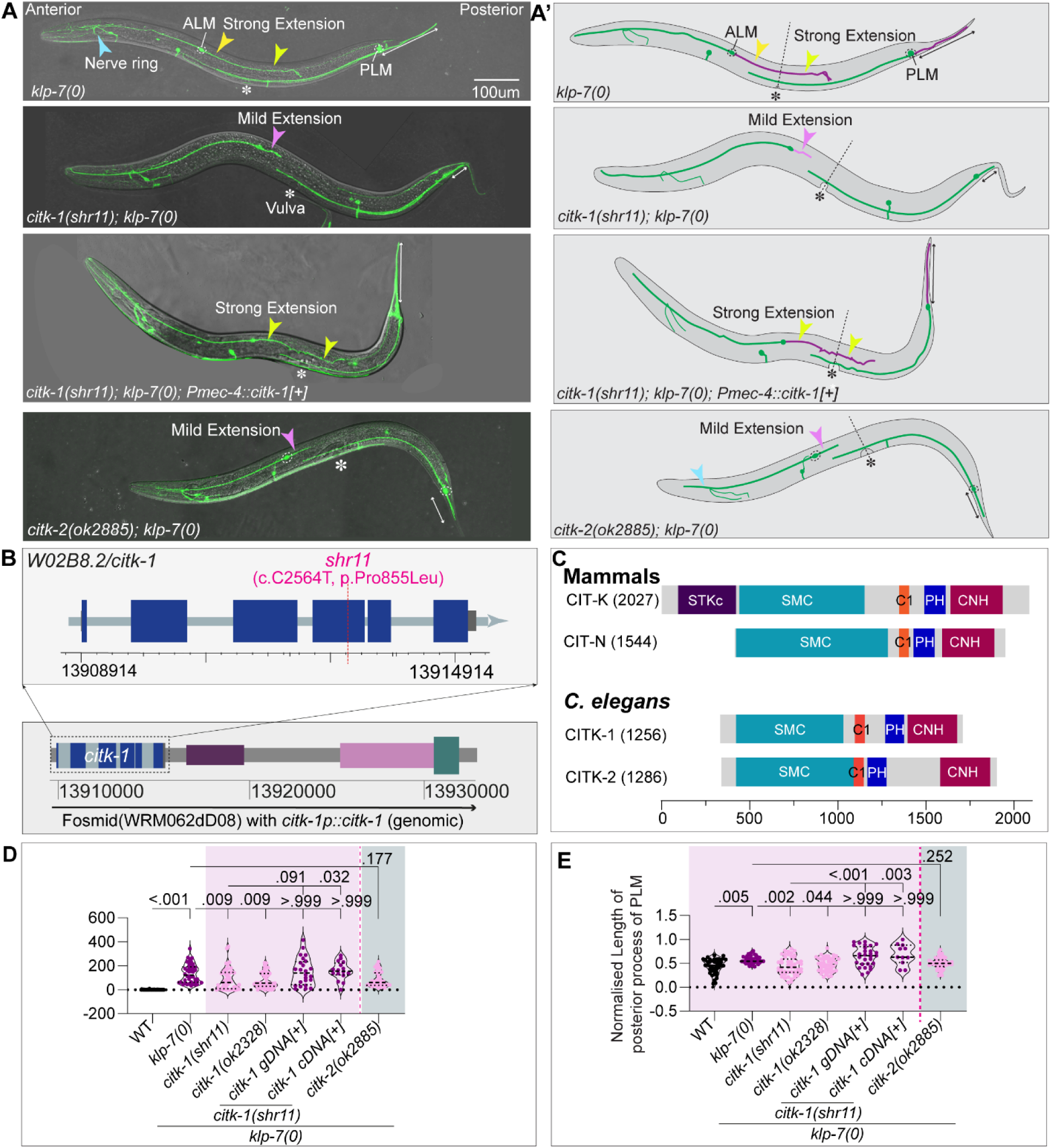
Identification and characterization of CITK-1 and CITK-2 as partial suppressors of *klp-7(0)* phenotype. (A-A’) Representative confocal images of the ALM and PLM mechanosensory neurons in the L4 stage *klp-7(0), citk-1(shr11); klp-7(0), citk-1(shr11); klp-7(0); shrEx432(Pmec-4::citk-1*[+]) and *citk-2(ok2885); klp-7(0)* mutant animals along with their respective schematics. These neurons are labeled green by *Pmec-7::GFP* reporter. The yellow-arrowheads point to the strong ectopic extension from the ALM neuron that crosses the vulval landmark, labeled by an asterisk (*) in *klp-7(0)* animals and the *citk-1(0); klp-7(0)* animals with an extrachromosomal expression of *Pmec-4::citk-1*. *Pmec-4* is a mechanosensory neuron specific promoter. The pink arrowheads point to the mild ectopic extension in *citk-1(shr11); klp-7(0)* and *citk-2(ok2885); klp-7(0)*. The double-headed white arrow extends along the length of PLM posterior process. (B) The schematic of the exon and intron of the *W02B8.2/citk-1* gene and the genomic positions of the identified candidate mutation within the *citk-1* locus of the *shr11* suppressor. At the bottom is a representation of the fosmid WRM062dD08 which contains the complete genomic cassette of the *citk-1* gene. (C) The schematics of the two isoforms of mammalian citron-rho interacting kinase proteins and the *C. elegans* paralogs CITK-1 and CITK-2. Adapted from (44-46). (D-E) Quantification of the absolute length of ALM ectopic extension (D) and normalized length of the posterior process of PLM neuron (E) in the WT, *klp-7(0), citk-1(shr11); klp-7(0), citk-1(ok2328); klp-7(0), citk-1(shr11); klp-7(0); citk-1* genomic DNA [+], *citk-1*(shr11); *klp-7(0); citk-1* cDNA [+], *citk-2(ok2885); klp-7(0)* backgrounds, where genomic *citk-1* is the *citk-1* fosmid WRM062dD08 and the *citk-1(cDNA)* is expressed under promoter *Pmec-4*. For D and E, N = 3 independent replicates, n (number of neurons) = 16-50. Normalized length of PLM posterior = (Absolute length/distance between the PLM cell body to the tip of the tail for the posterior neurite). For D-E, Error bars represent SEM (Standard error mean), P values from Kruskal Wallis test followed by Dunn’s multiple comparisons.

Several lines of evidence confirmed that *shr11* is an allele of *W02B8.2*. First, the mapping results were validated by an extrachromosomal expression of a wild-type copy of *W02B8.2* gene in the *klp-7(0); shr11* suppressor background (fosmid WRM062dD08, which contains the complete genomic cassette of *W02B8.2* gene) (Figure 2B). This expression rescued the ALM ectopic extension phenotype of the *klp-7(0)* mutant (Figure 2A-A’, D, S2B) which confirmed that *W02B8.2*(*shr11)* missense mutation partially suppresses the *klp-7(0)* phenotypes. Next, the partial suppression phenotype was reproduced by introducing a deletion allele of *W02B8.2*, *ok2328* (Figure S2G), in *klp-7(0)* mutant background. Furthermore, an extrachromosomal expression of *W02B8.2* cDNA under a Touch-receptor neuron specific promoter, *Pmec-4*, could also rescue the ectopic extension phenotype of *klp-7(0)* in *W02B8.2(shr11); klp-7(0)* suppressor background (Figure 2A-A’, D, S2B) implying a cell-autonomous role of *W02B8.2* in suppression of *klp-7(0)* phenotype.

Further, sequence and structural blast analysis confirmed that W02B8.2 is a kinase-less orthologue of mammalian citron-rho-interacting kinase and has another paralog in worms, F59A6.5, with which it shares 66% sequence homology (Figure S2C-D). Using the protein databases (44-46), we identified that similar to the neuronal isoform of mammalian CIT (CIT-N) which lacks an N-terminal kinase domain (47), W02B8.2 and F59A6.5 share an extended coiled-coil domain, a phorbol ester/DAG type Zn finger domain (C1 domain), a pleckstrin homology (PH) domain, and a Citron-Nik-1 homology (CNH) domain with the mammalian citron kinase protein (Figure 2C, S2E-F) and hence we named these loci *citk-*1 and *citk-2,* respectively.

The identification of a paralog *F59A6.5/citk-2* and observed partial but not complete suppression of *klp-7(0)* phenotype by both the alleles of *W02B8.2/citk-1, shr11* as well as *ok2328,* suggested a functional redundancy. Therefore, we assessed if *citk-2* mutation suppresses the ALM ectopic extension phenotype of *klp-7(0)* mutant, phenocopying *citk-1(0).* We used a frameshift deletion variant of *citk-2, ok2885*, to evaluate the suppression (Figure S2H). A double mutant of *citk-2(ok2885); klp-7(0)* displayed significantly shorter ALM ectopic extension than the *klp-7(0)* mutant, suggesting a partial suppression of *klp-7(0)* phenotype (Figure 2A-A’, D, S2B)).

As previously reported *klp-7(0)* mutant animals also display an overgrowth of the posterior process of PLM neurons (2) (Figure 2A-A’, E). The PLM neurons are the second pair of bilateral mechanosensory neurons running ventrolaterally along the ventral nerve cord in the posterior half of the worm. Unlike ALM neurons, PLM neurons are bipolar with a long axon-like anterior process and a short dendrite-like posterior process that extends into the tail of animal (Figure 2A-A’). We observed that the second-site mutations in *citk-1* locus also suppresses the PLM posterior overgrowth phenotype of *klp-7(0),* which could be rescued by extrachromosomal expression of WT copy of either genomic or cDNA of *citk-1* (Figure 2A-A’, E). The *citk-2(0); klp-7(0)* double mutant however shows a decreasing trend but not a significant suppression of the PLM posterior overgrowth phenotype of *klp-7(0)* mutant, reiterating our redundancy hypothesis.

In conclusion, mutations in the kinase-less worm orthologues of mammalian citron-kinase protein, *citk-1* and *citk-2,* can partially suppress the microtubule-dynamics-associated neuronal ectopic extension phenotypes of *klp-7(0)* mutant.

### CITK proteins regulate dynamic microtubules in the anterior process of PLM neurons and microtubule arrangement in the posterior process of PLM neurons

Building on our previous observations we investigated the effects of loss of *citk-1* and *citk-2* on microtubule dynamics in the PLM mechanosensory neurons. We conducted time-lapse live-imaging of a fluorescently tagged reporter, EBP-2::GFP, to track the growing plus-ends of microtubules (2, 22, 25). As previously described (2), analysis of kymographs generated from 30 µm regions of interest spanning anterior and posterior processes of PLM neurons can be used to determine the microtubule polarity and dynamics in these neurons (Figure 3A).

**Figure 3.**
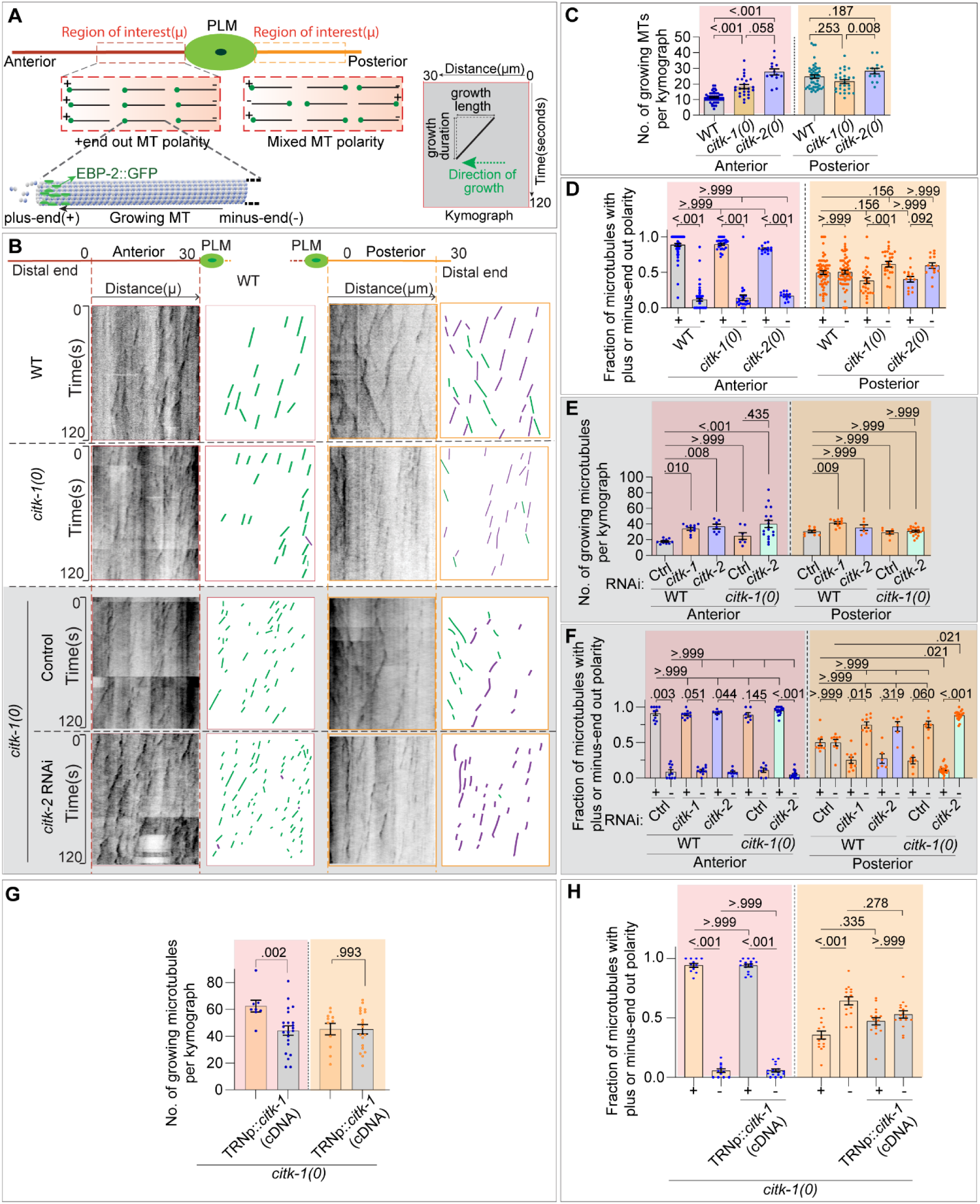
CITK-1 and CITK-2 synergistically regulate microtubule dynamics in PLM neurons. (A) Schematic representation of the PLM anterior and posterior processes, with red and orange dotted rectangles delineating the regions of interest (ROIs) utilized for analyzing time-lapse live images of *Pmec-4*::EBP-2::GFP (*juIs338*). Inset is a pictorial representation of a kymograph (distance versus time plot) that illustrates the movement of EBP-2::GFP along the growing microtubule in a neuronal process. The slope of the track in the kymograph is a ratio of growth length and growth duration of the polymerizing microtubule, with the direction of the slope indicating the direction of growth. **(B)** Representative kymographs and schematics of the EBP-2::GFP reporter obtained from the live-imaging of aforementioned ROIs in the wild-type, *citk-1(0),* and *citk-2(0)* (white background), and RNAi strain (in gray background) *citk-1(0); lin-15b (-); shrsI2(Pmec-3-sid-1+unc-119*[+]) fed on control and *citk-2 RNAi* bacteria. *Pmec-3-sid-1* is used to overexpress *sid-1* and enhance RNAi sensitivity in tissues expressing *Pmec-*3, which includes PLM mechanosensory neurons. In the schematics, green and purple traces represent movements of EBP-2 bound microtubules away from the cell body (Plus end out) and movements towards the cell body (Minus end out), respectively. **(C-D)** Quantification of the number of observed diagonal tracks of EBP::GFP/growing microtubules (C) and fractional polarity (the fraction of microtubules with ‘plus-end-out’ (+) or ‘minus-end out’ (-) polarity) (D) in the two processes of PLM neurons in the wild-type, *citk-1(ok2328)* and *citk-2(ok2885)* mutant backgrounds. N = 3–4 independent replicates, n (number of neurons) = 14-63. **(E-F)** Quantification of the number of growing microtubules (E) and fractional polarity (F) in the anterior and posterior processes of PLM neurons in the wild-type control and *citk-1(0)* strains with a *lin-15b(-); shrsI2(Pmec-3-sid-1+unc-119*[+]) background for enhanced RNAi sensitivity. For RNAi mediated knockdown the wild-type animals and *citk-1(0)* animals were fed on bacteria with an empty vector (control) or an expression cassette for *citk-1* or *citk-2* dsRNA. N = 3 independent replicates, n (number of neurons) = 7-20. **(G-H)** Quantification of the number of growing microtubules (G) and fractional polarity (H) in *citk-1(0)* mutant animals and the *citk-1(0); shrEx512(Pmec-4::citk-1(cDNA)*[+]) animals with an extrachromosomal copy of *citk-1(cDNA)* expressed under *Pmec-4.* N = 3 independent replicates, n (number of neurons) = 8-22. For C-F, H, Error bars represent SEM (Standard error mean), P values from Kruskal Wallis test followed by Dunn’s multiple comparisons. For E, Error bars represent SEM (Standard error mean), P values from Mann-Whitney test. The groups separated by dotted lines were analyzed independent of each other.

We observed that in comparison to wild-type animals, both *citk-1*(*0*) and *citk-2(0)* mutant animals displayed an increase in the number of growing microtubules/EBP-2::GFP comets in the PLM anterior process (Figure 3B-C), accompanied by decreased growth length and growth duration of these microtubule growth/polymerization events (Figure 3B, S3A-B), suggesting an increase in the plus end microtubule dynamics in the PLM anterior process. In contrast, the posterior process did not show significant change in the number of growing comets (Figure 3B-C, S3A-B). However, the growth length and growth duration of these microtubule growth/polymerization events showed an increase in *citk-1(0)* animals (Figure 3B-C, S3A-B) suggesting decreased microtubule dynamics in the posterior process.

Furthermore, the arrangement of microtubules in the anterior and posterior processes of PLM neurons is well defined (2, 25, 48). The anterior process is known to show plus-end-out microtubule polarity similar to the vertebrate axons and accordingly, the kymographs display a majority of diagonal tracks directed away from the cell body (plus-end-out, green trace) (Figure 3A-B, D). In contrast, the posterior process is known to display a mixed microtubule polarity, similar to the vertebrate dendrites and hence, kymographs exhibit a comparable fraction of comets directed away from (plus-end-out, green trace) and towards the cell body (minus-end-out, purple trace) (Figure 3A-B, D). We observed that the mutations in neither *citk-(0)* nor *citk-2(0)* perturb microtubule polarity in the anterior process of PLM neurons (Figure 3B, D). However, the number of minus-end-out tracks in the posterior processes of PLM neurons of *citk* mutant animals seemed to be significantly higher than the number of plus-end-out comets implying a trend towards minus-end-out polarity (Figure 3B, D). The observed similarities in the mutant phenotypes along with the structural and molecular homology reiterated a degree of redundancy in the functions of *citk-1* and *citk-2*. Therefore, we assessed the impact of loss of both CITK-1 and CITK-2 on microtubule dynamics in the PLM neuron. We first tried generating *citk-1(0)* and *citk-2(0)* double mutants, however, we could not generate a viable double homozygote. Additionally, two previous reports have shown that the knockdown of both *citk-1/W02B8.2* and *citk-2/F59A6.5* leads to gonadal sterility and embryonic lethality (43, 49). So, we focused on a neuron-specific knockdown of *citk-1* and *citk-2,* by double-stranded RNAi using a feeding method (50). However, simultaneous feeding of two different RNAi bacteria has been shown to reduce RNAi efficiency (51–53). Therefore, we knocked down *citk-2,* in the *ok2328* deletion mutant of *citk-1,* by feeding method in an RNAi-sensitive strain, with enhanced sensitivity in mechanosensory neurons (54). We observed that single knockdown of either *citk-1* or *citk-2* in the RNAi sensitive strain displayed microtubule dynamics and polarity defects similar to those observed in single mutant animals (Figure3B, E-F, S3C-D). The strength of the defects observed in the single *citk-1* mutant animal was further aggravated by knockdown of *citk-2* in the *citk-1* mutant background. The number of microtubules growing in the anterior process showed a further increase than the single mutant animals (Figure 3B, E). Additionally, the posterior process of PLM neurons revealed a significant increase in the fraction of minus-end-out comets compared to single mutant animals, suggesting a minus-end-out microtubule polarity, even though the plus-end-out microtubule polarity of anterior process remained unaffected, (Figure 3B, F). Moreover, the growth length and growth duration additively decreased in the PLM anterior process suggesting a synergistic increase in the MT dynamics on loss of both *citk-1* and *citk-2* (Figure 3B, S3C-D). Moreover, the PLM posterior processes displayed an increased growth length and growth duration on knock-down of *citk-2* in wild-type as well as *citk-1(0)* animals suggesting a decreased microtubule dynamics in PLM posterior process, however, the effect is not additive (Figure S3C-D).

We further validated the cell-autonomous role of *citk-1* by expressing a wild-type copy of *citk-1*(cDNA) under touch receptor neuron specific promoter in the *citk-1(0)* mutant animals. The number of growing microtubules in the anterior process of PLM neurons in the rescued worms is less than the mutant animals (Figure 3G), while the growth length and growth duration increases (Figure S3E-F). Moreover, the rescued animals also show a mixed microtubule polarity unlike the trend towards minus-end-out MT polarity in the mutant animals (Figure 3H). Thus, the defects in microtubule dynamics and polarity are rescued by the extrachromosomal copy of wild type *citk-1*(cDNA) suggesting a cell autonomous role of *citk-1* in regulating microtubule dynamics and polarity.

These observations suggest that *citk-1* and *citk-2* redundantly rein in the number of dynamic microtubules in the anterior and posterior process of PLM neurons and maintain the plus-end-out population of microtubules in the posterior process of PLM neurons, for a mixed microtubule polarity in the posterior process.

### PLM mechanosensory neurons displayed growth and guidance defects on loss of both *citk-1* and *citk-2*

As previously described, perturbations in the microtubule dynamics are associated with changes in the structure and function of these neurons, for example observed ectopic growth in the *klp-7(0)* animals (2, 30, 42). Therefore, to investigate the effects of the observed microtubule perturbations on the neuronal structure in *citk-1* and *citk-2* mutant animals we assessed the morphology of bipolar PLM mechanosensory neurons in larval stage 4 animals. As previously described, the wild-type PLM neurons extend a long anterior process that extends towards the midbody running ventrolaterally parallel to the ventral nerve cord past the vulva (marked by an asterisk) (Figure 4A, S4A-A’). In our analysis, we observed no significant alterations in the morphology of PLM neurons in *citk-1* or *citk-2* single mutants (Figure S4B-D). Both, the loss of function mutant, *citk-1(ok2328),* as well as the missense mutant, *citk-1(shr11),* displayed a significantly shorter PLM posterior process compared to the wild-type animals (Figure S4A-B). However, the length of the PLM anterior process in *citk-1* mutants was similar to the wild-type animals (Figure S4C). Occasionally the PLM anterior process would wander off to the dorsal side, suggesting a guidance or migration defect (Figure S4A-A’). Similar to *citk-1(0)* mutant animals the *citk-2(0)* mutant did not display any significantly drastic defect in the morphology of PLM neuron apart from rare migration defect (Figure 3.3A, C). Furthermore, the ventral synaptic branch, which forms a little posterior to the vulva in wild-type animals (marked by a white arrow in Figure 3.3A), got misplaced to ectopic positions or there are additional ectopic synapses in 11% of *citk-1(ok2328),* 7% of *citk-1(shr11)* and 6% of *citk-2(ok2885)* mutant animals. (Figure 3.3E, F).

**Figure 4.**
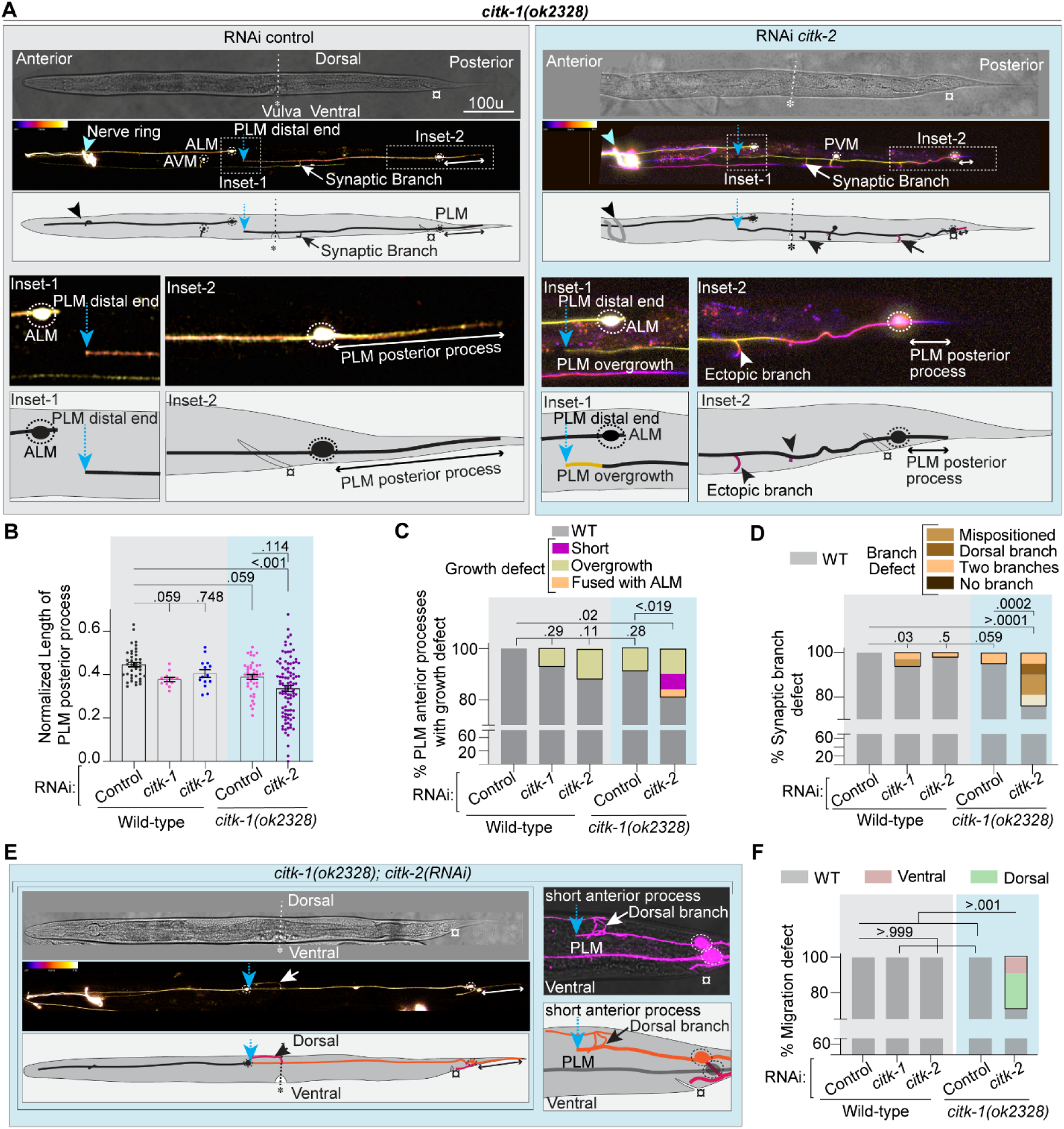
Growth and guidance defects observed on knockdown of *citk-2* in *citk-1(0)* animals. **(A**) Representative confocal images and schematics of ALM and PLM mechanosensory neurons of L4-stage *citk-1(0)* mutant animals with an RNAi sensitive background *lin-15b(-); shrsI2(Pmec-3-sid-1+unc-119*[+]) exposed to either control or *citk-2* RNAi feeding. The head of the worm (anterior end) is placed towards left, the tail (posterior end) points to the right, the dorsal edge of its body faces upward and the ventral edge with vulva faces downwards in the images. The confocal image is split into DIC channel image (top panel) and Z-axis temporally profiled fluorescence intensities (lower panel) image with respective color coding maps placed on the top-left corner. The mechanosensory neurons in these animals are labeled by a *Pmec-4::mCherry (tbIs222)* reporter. Inset-1 zooms in on the distal end (marked by a blue dotted arrow) of anterior process of PLM neuron and the location of ALM cell body. Inset-2 focuses on the extent of PLM posterior process (marked by a double-headed white arrow). The white arrow points to the ventral synaptic branch and the white arrowhead marks the ectopic synaptic branch. Vulva is marked by an asterisk (*), anus is marked by a generic symbol (¤) and the celeste-blue arrowhead points to the nerve ring. **(B)** Quantification of the normalized length of PLM posterior process in the wild-type control and *citk-1(0)* RNAi sensitive background animals fed on bacteria expressing either empty vector (control) or *citk-1* or *citk-2 ds* RNA for RNAi mediated knockdown. **(C)** Plot of percentage of PLM neurons exhibiting anterior neurite growth defects, PLM anterior neurite ends before vulval position: short neurite defect (magenta), PLM anterior neurite extending past ALM cell body: Overgrowth defect (pale-green), PLM anterior neurite is misguided or grows and merges with the Z planes of ALM neuron, appears to be fused with ALM (pale orange). The percentage of animals with wild-type like morphology is depicted in gray. **(D)** Quantification of percentage of PLM neurons displaying a ventral synaptic branch defect, which in wild-type animals is localized a little posterior to vulva (labeled by white arrow in (A). In mutant animals it can be mispositioned (light-brown) to an alternate position along the length of PLM neuron, can be directed towards dorsal side of animal: Dorsal branching (brown), can be over-represented: Two branches (cream) or can be completely absent (dark-brown). B-D, N = 3 independent replicates, n (number of neurons) = 14-96. **(E)** Representative confocal images and schematics of dorsal mispositioning and neurite growth defects seen in *citk-1(0)* mutant RNAi sensitive animals fed for RNAi mediated knockdown of *citk-*2. The worm is placed in the same manner as control worms in panel (A). The blue dotted arrow marks the distal tip of the anterior process of PLM neurons and the white double-headed arrow labels the PLM posterior process. The white arrow points to the dorsally directed synaptic branches. Asterisk symbol (*) labels ventrally positioned vulva and the generic symbol (¤) marks the anal position. **(F)** Quantification of the percentage of animals exhibiting a guidance defect. One of the two PLM neurons cell body in these animals is either mispositioned along the dorsal edge of body: Dorsal (green), or along the ventral nerve cord(VNC): ventral (dusty pink). The wild-type animals have a ventro-lateral placement of PLM cell body growing parallel to the VNC: Wild-type(gray). N = 3 independent replicates, n (number of animals) = 12-96. For B, Error bars represent SEM (Standard error mean), P values from Kruskal Wallis test followed by Dunn’s multiple comparison test. For C-D, P values from Fisher’s exact test for WT phenotype and defect phenotype.

Since our previous observations suggested a redundant role of both citron kinases in microtubule regulation, therefore we studied the effects of knockdown of *citk-2* in *citk-1(ok2328)* mutant animals (as described in previous section) on the morphology of PLM neurons to understand their role in neuron development. We observed a spectrum of weakly penetrant growth and guidance defects in the PLM mechanosensory neurons of these animals.

As shown in Figures 4A-B, the *citk-1(0)* single mutant animals display a visible shortening of posterior processes of PLM neurons, which is synergistically further reduced on loss of *citk-2* in *citk-1(0)* mutant animals (Figure 4A, B). In contrast the anterior processes of most of the PLM neurons (80 out of 97) on *citk-2* knock down (kd) in *citk-1(0)* mutant animals displayed wild-type like growth, where PLM neuron terminates near or before the ALM cell body, similar to the single mutant animals (Figure 4A, S4A-A’). However, 9 of the 97 PLM neurons analyzed displayed an “overgrowth phenotype” characterized by its overshooting past the ALM cell body (Figure 4A, C) and 6 of the 97 PLM neurons displayed a “short neurite defect”, characterized by premature termination of the anterior process before vulval position (Figure 4C, E), while the distal end of 3 of the PLM anterior processes could not be identified as it was indistinguishable from the ALM cell body due to the guidance defects (Figure 4C, E). However, quantification of the lengths of PLM anterior processes did not show any significant difference from the wild-type animals (Figure S4E).

Interestingly, the growth and guidance defects observed correlated with the mispositioning of the PLM neuron cell body (Figure 4E). The PLM cell body in wild-type animals lies ventrolaterally, a little behind the anus and extends a long anterior process that grows ventrolaterally parallel to the ventral nerve cord (VNC) and a short posterior process that extends into tail along its ventral edge (Figure S4A). In the *citk-1(0); citk-2(kd)* animals one of the two PLM neurons in twenty-seven out of 94 animals was mispositioned, either dorsally, towards the dorsal nerve cord (19 in 94 animals), or ventrally, juxtapositioned to the ventral nerve cord (8 in 94 animals) (Figure 4E-G). As one would expect, the mispositioning of these PLM neurons was accompanied by either growth defect like short neurite defect or guidance defect where the anterior process of PLM neurons extends dorsolaterally along the dorsal side or ventrally juxtapositioned to ventral nerve cord or finds its way back to its ventrolateral path (Figure 4E-F, S4F).

Furthermore, we observed that growth and guidance defects led to irregularities in effective synaptic branch formation. (Figure 4D). The dorsally mislocalized PLM neurons extend dorsolaterally and therefore protract branches towards the dorsal nerve cord, failing to form an effective ventral synaptic branch (Figure 4D-E, S4F). The anterior processes of ventrally mislocalized PLM neurons are juxtapositioned to the ventral nerve cord and hence its effective synaptic partner in VNC, with no clear demarcation of a synapse (Figure S4F). While in remaining cases with a mispositioned PLM neuron the position of the synaptic branch may vary, or the synaptic branch may be altogether absent (Figure 4D). Additionally, we observed ectopic protrusions emanating from the PLM cell body and anterior as well as posterior processes in *citk-1(0); and citk-2* (RNAi kd) animals (Figure S4G).

These observed structural defects such as ectopic protrusions and growth defects are markers of altered microtubule dynamics in the PLM neurons (30, 33), reiterating the roles of *citk-1* and *citk-2* in regulating the microtubule dynamics and hence, growth and guidance of PLM mechanosensory neurons. However, the precise mechanism by which citron kinase regulates microtubule dynamics and hence, neuron structure and maintenance remains unclear.

### Microcephaly associated protein ASPM-1 phenocopies the loss of citron kinase proteins CITK-1 and CITK-2

Previous molecular studies have shown that mammalian citron kinase is recruited by ASPM, a microtubule minus-end associated protein, to spindle poles during cytokinesis to regulate organization of spindle microtubule arrays (55, 56). Additionally, both proteins have been shown to colocalize to the midbody region during cytokinesis (57). Similarly, the worm orthologue of ASPM, ASPM-1, localizes to the spindle poles and is required for spindle pole organization. Protein network analysis (58) for interactors of *citk-1* and *citk-2* also showed strong interactions with *aspm-1* (Figure 5A). Drawing on these findings, we hypothesized that citron kinase interacts with ASPM-1 in neurons to regulate microtubule dynamics.

**Figure 5.**
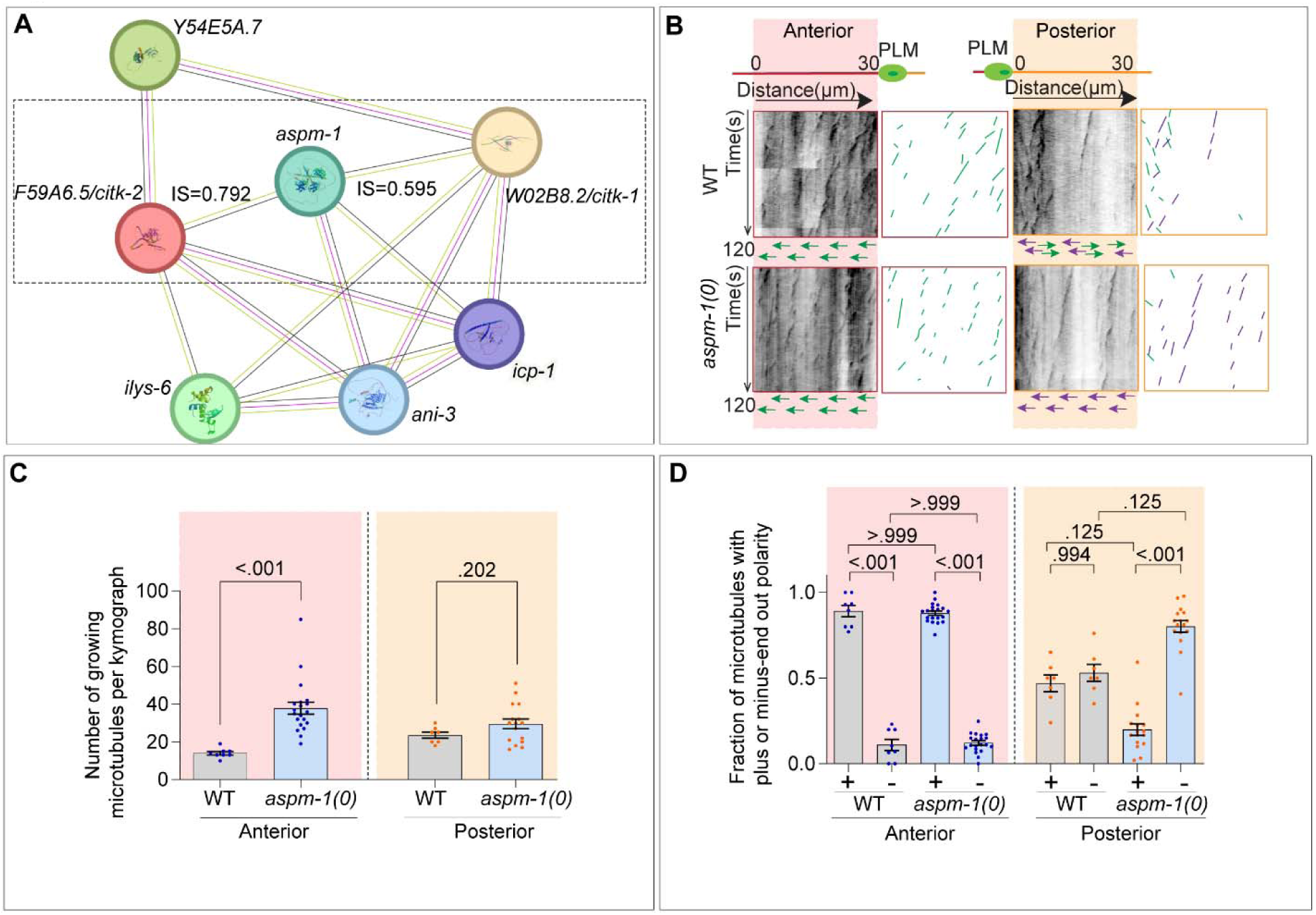
Microtubule dynamics and fractional polarity defects observed in *aspm-1(ts)* mutants. **(A)** Protein-protein interaction network obtained for *citk-1* and *citk-2* using STRING database. The nodes represent query and known interacting proteins. Edges represent known and predicted protein-protein functional associations (not necessarily physical interaction) between these proteins or their putative homologs in varied species. The type of association is color coded based on evidence for association such as: Magenta edges (°—°): experimentally determined association based on biochemical or genetic data (including experimental evidences for interaction of putative homologs in other species), Black edges (°—°): co-expression, Green edges (°—°): text mining (Co-mentioned in PubMed abstracts). ‘IS’ is the combined interaction score which represents an approximation of confidence in a given association being true, based on all available evidence, on a scale of zero to one. An interaction score of 0.5 or less indicates 50% incidence of false positives. CITK-1 and CITK-2 have no known direct physical association, however, both show interaction with ASPM-1 with a confidence score >0.5 **(B)** Representative kymographs and schematics obtained by plotting the movement of EBP-2::GFP (*juIs338)* comets in the anterior and posterior processes of PLM neurons (from ROIs mentioned in Figure 3) of the wild-type and *aspm-1(0)* animals grown at non-permissive temperature of 25°C. (0) represents loss of function temperature sensitive allele of *aspm-1.* The diagonal lines represent the number of EBP-2::GFP tagged plus-end of microtubules growing away from (green) or towards (purple) the PLM cell body. The slope of the line is the ratio of the growth length and growth duration of a polymerization event. **(C-D)** Quantification of the number of growing microtubules/EBP-2::GFP comets (C) and the fraction of plus-end-out or minus-end out microtubules in the wild-type and *aspm-1(0)* mutant animals grown at non-permissive temperature. N=3 biological replicates, n (number of neurons) = 7-16. Error bars represent SEM (Standard error mean), For C, P values from Mann-Whitney test and for D, P values from Kruskal Wallis test followed by Dunn’s multiple comparison test. The groups separated by dotted lines were analyzed independent of each other.

We used a temperature sensitive loss of function variant of *aspm-1* (59), to characterize its role in regulation of microtubule dynamics in neurons as previous widescreen RNAi studies have associated *aspm-1* knockdown with embryonic sterility and embryonic lethality, similar to loss of both *citk-1* and *citk-2* (60). Using the EBP-2-GFP reporter (Figure 3A), as described in previous section, we conducted live imaging of the end-dynamics of growing microtubules and observed a significant increase in the number of growing microtubules (EBP-2::GFP comets) in the anterior process of PLM neuron similar to citron kinase mutants (Figure 5B-C). Similarly, the growth length and growth duration of these microtubule growth/polymerization events in the PLM anterior processes showed a significant reduction when compared to wild-type animals (Figure 5B, S5A-B). Furthermore, the PLM posterior processes displayed a significant increase in the growth length, however no change in the number of growth events or growth duration as in citron kinase mutants (Figure 5B-D, S5A-B). Additionally, although the PLM anterior processes did not show any change in the microtubule polarity, the PLM posterior process displayed a significant increase in the number of minus-end-out growing microtubules suggesting a minus end-out microtubule polarity in *aspm-1(ts)* mutant animal (Figure 5C, E), similar to citron kinase mutants.

Building on this observation that *aspm-1* mutation increases microtubule dynamics in the PLM mechanosensory neurons, similar to citron kinase mutants, we further evaluated if it would be able to suppress the ectopic extension phenotype of *klp-7*(*0*) mutant animals linked with reduced microtubule dynamics. And as expected, the introduction of *aspm-1(ts)* mutation in the *klp-7(0)* mutant background could partially suppress the ALM ectopic extension phenotype (Figure S5C-D). However, the overgrowth phenotype of PLM posterior was not suppressed.

These observations suggested that ASPM-1, similar to citron kinase differentially regulates microtubules in the anterior and posterior processes of PLM neurons. ASPM-1 regulates microtubule stability in the anterior process and plus-end-out microtubules in the posterior process. And *aspm-1* and citron kinases work antagonistically to KLP-7 in regulation of microtubule dynamics in the ALM mechanosensory neurons.

## Discussion

In this study, we conducted a genetic screen to identify novel regulators of microtubule dynamics and isolated a mutation in W*02B8.2/citk-1,* a kinase-less worm orthologue of mammalian citron-rho interacting kinase (CIT), which has another kinase-less paralog *F59A6.5*/*citk-2* in worms. Our findings demonstrated that *citk-1* and *citk-2* have redundant roles in microtubule stabilization in the axon-like anterior process and organization of microtubule polarity in the dendrite-like posterior process, and hence growth and guidance of PLM neuron. Further findings demonstrate that citron kinases work in the same genetic pathway as *aspm-1,* a microtubule minus-end associated protein, to regulate microtubule dynamics in PLM neurons (Figure 6).

**Figure 6.**
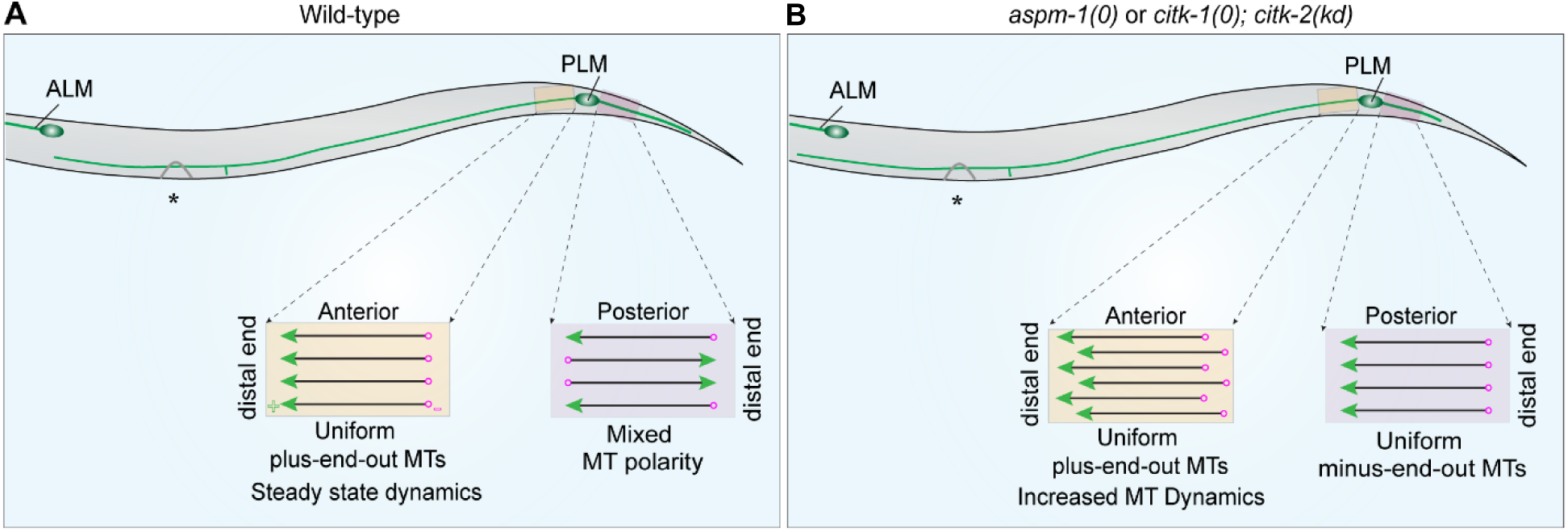
Proposed model. Schematic representation of microtubule dynamics in the anterior and posterior processes of PLM mechanosensory neurons in the wild-type *C. elegans* animals (left panel) and the mutant animals (right panel). The 30 µm ROI in the anterior process (orange rectangular selection) and posterior process (pink rectangular selection) are zoomed-in to illustrate microtubule dynamics in these processes. The black lines represent microtubules, the green arrowhead depicts the plus-ends of microtubules, bound by plus-end-binding proteins and the circular ring depicts the minus-ends of microtubules. The direction of arrowhead shows the direction of microtubule growth. **(A)** In the wild-type animals the anterior process exhibits a plus-end-out steady-state microtubule dynamics, while the posterior process displays a mixed microtubule polarity. **(B)** Similar ROI in mutant animals shows increased microtubule dynamics in the anterior process, schematized as increased number of growing microtubules, and a minus-end-out microtubule polarity in the posterior process.

### CITK and ASPM-1 maintains axonal microtubule stability in PLM neurons

Citron-rho interacting kinase was originally identified as a Rho/Rock effector and was later described to be involved in regulation of cytokinesis through organization of spindle microtubule arrays and midbody maturation (61). A previous study suggested that the role of CIT proteins in cytokinesis is conserved and the two orthologues of CIT in worms, *W02B8.2/citk-1* and *F59A6.5/citk-2* function redundantly (43). Using available protein databases and literature (44–46), we found that these two proteins share a sequence and structural homology, as well as functional homology. We also found that these two proteins are structural homologs (62, 63) of neuronal isoform of mammalian CIT that lacks an N-terminal kinase domain. Furthermore, it has been shown that CIT-K regulates neurite extension and dendrite morphology *in-vitro* and *in-vivo*, while limiting axon outgrowth in regenerating DRGN neurons *in-vitro* (64, 65). Our data aligns with these observations, as loss of either or both citron kinases results in shortening of dendrite-like posterior process of PLM neurons, while the axon-like anterior process displayed an overgrowth phenotype on loss of both citron kinases. Previously, it was suggested that CIT-K regulation of neurite morphology overlaps with modulation of actin cytoskeletal dynamics, however, the mechanistic details of this regulation were unclear (64–67). Another study showed that in dividing HEK cells, citron kinase regulates the length, stability and nucleation of astral microtubules (56).

Interestingly, in this study, we found that the kinase-less, citron kinase regulates the population of dynamic microtubules in the anterior processes of PLM neurons without any effect on microtubule plus-end-out polarity, suggesting that citron kinase reigns in the number of dynamic microtubules in the axon-like processes, similar to other subtle regulators of microtubule dynamics in PLM neurons such as PTRN-1/CAMSAP (2, 32, 33). However, the molecular basis of how CITK regulates microtubule stability in the PLM anterior process is not clear.

The CITK protein through its interactions with Rho GTPase pathway is enriched in the midbody region during cytokinesis (68–71). Additionally, along with its interacting partners such as Centralspindilin complex and KIF14, CITK regulates cross-linking and bundling of microtubules in midbody (72–75). Thus, whether CITK retains a similar role in cross-linking the parallel bundles of axonal microtubules, analogous to midbody, needs further investigation. Moreover, Myosin-II-RhoA complex is part of the cortical-actin in axon, which plays an important role in regulating axonal morphology and rigidity (76–78). This suggests that the protein complex that bridges the actin and microtubule cytoskeleton during cytokinesis could also be at the interface of regulation of actin dynamics and microtubule dynamics in neuron development, similar to other microtubule associated proteins such as kinesin Kif21b that apart from its role in regulation of microtubule dynamics also modulates actin cytoskeleton to regulate radial migration of cortical projection neurons (79, 80). And interestingly enough, we show that citron kinase mutant animals also display cell positioning and guidance defects in PLM mechanosensory neurons.

It is worth mentioning that mutations in mammalian citron kinase were associated with MCPH (autosomal recessive primary microcephaly), a small brain disorder linked with mild to severe delayed cognitive development and behavioral challenges, in last decade (81, 82). From studies in CITK-/-mice these defects were attributed to defects in division of neuronal precursor cells, apoptosis and cell migration (61, 83). In this study we found that loss of citron kinase leads to perturbation in microtubule dynamics and neurite growth in PLM neurons. Therefore, it is worth testing if perturbations in microtubule dynamics in differentiated neurons could also be a contributing factor.

Interestingly, more than 40% of MCPH associated variants are found in the locus of ASPM, a microtubule minus-end associated protein, which was previously shown to recruit citron kinase to spindle poles in dividing HEK cells (56, 84, 85). It was shown that citron kinase is a crucial downstream partner of ASPM in regulation of dynamics of microtubule spindle arrays (56). Previous studies have demonstrated that ASPM-1 plays a crucial role in meiotic spindle rotation in oocytes in *C. elegans* which is evolutionarily conserved in higher mammals as well (59, 84, 86, 87). In mammals, ASPM is expressed in neuronal precursor cells and regulates neurogenesis by directing spindle axis rotation and symmetric divisions in neuronal precursor cells, neuronal migration and cell fate acquisition (86, 88–90). However, the role of ASPM in differentiated neurons has not been characterized.

Interestingly, here we show that similar to citron kinase, ASPM-1 limits the number of dynamic microtubules in the axon-like anterior process of PLM mechanosensory neurons similar to citron kinase, and PTRN-1 (32, 33, 91). Thus, suggesting that ASPM-1 also has post-mitotic roles in PLM neurons. Recent reports have also shown that cytokinesis associated proteins are repurposed in differentiated neurons (92).

### CITK and ASPM-1 maintain the plus-end-out microtubule population in the dendrite like posterior neurite of PLM neuron

ASPM, similar to CAMSAP, has been shown to regulate minus-end dynamics of microtubules *in-vitro* (21, 55, 93). In this study we showed that similar to citron kinase, ASPM-1, the worm orthologue of ASPM, apart from its role in cytokinesis, regulates the population of plus-end-out microtubules in the dendrite-like posterior process of microtubules (Figure-6). This is in sharp contrast to PTRN-1/CAMSAP, which regulates the minus-end-out population in the posterior process of PLM neurons (2).

It has been shown previously that CIT-K works downstream of ASPM in regulating spindle arrays, and ASPM and CIT-K also colocalize in the midbody region as well (56, 57). Our genetic analysis also suggests that ASPM and CIT-K work in the same genetic pathway to regulate PLM neuron morphology. It will be interesting to explore if citron kinase is being recruited by ASPM to the minus-ends of microtubules in PLM neurons. Furthermore, how ASPM-1 and CITK maintain the plus end out MT population in the posterior PLM neurite is not clear. Whether ASPM-1/CITK complex competes with PTRN-1 to limit minus-end-out population, or they have independent roles in plus-end-out growth of microtubules needs to be explored.

In summary, in this study we show two microcephaly associated proteins involved in cytokinesis, Citron Kinase and ASPM-1, which are repurposed in post-mitotic PLM mechanosensory neurons to regulate microtubule dynamics and arrangement, along with the regulation of growth, guidance as well as positioning of these neurons.

## Materials and Methods

### C. elegans genetics

*C.* elegans strains were reared on the standard NGM (nematode growth medium plates) seeded with *E. coli* OP50 bacterial strain. For all experimental purposes worms were grown at 20°C, except for the temperature-sensitive(ts) strains (94) which were grown at non-permissive temperature 25°C according to experimental paradigms. The (0) denotes the loss of function allele, for instance, the *tm2143* deletion allele of *klp-7* has been at places mentioned as *klp-7(0)*. The strains studied in this paper, including the mutants generated by EMS mutagenesis, have been listed in Table S4. The published transgenes used in this study have also been listed in Table S4. The newly generated transgenic strains carrying extrachromosomal arrays of DNA are tabulated in Table S5. The transgenic lines were generated by microinjection into the gonad of the parent (95). The homozygosity of mutants was confirmed either by PCR genotyping or by sequencing.

### EMS mutagenesis

The suppressor mutants were generated by EMS mutagenesis (94). The age-synchronized larval-4 stage animals were pooled and washed thrice with the 1xM9 solution (isotonic salt solution), to get rid of any gut bacteria, following which, the animals were incubated in the 1xM9 solution containing 47mM EMS at 20°C for 4 hours on a hula mixer, which allowed continuous mixing and aeration of media. Post-incubation, the animals were pelleted, and the supernatant was discarded, followed by three washes with 1XM9 solution. Post-wash animals were pelleted, and 6-7 healthy animals were transferred to fresh NGM plates seeded with OP50 bacteria. The resulting F1s were transferred to fresh plates and the F2 generation animals were screened for suppression phenotype. Only one suppressor was picked from one F1 plate. Interestingly, in all cases, the suppression phenotype in one F1 plate was shown by about 25% of the F2 animals, suggesting that the EMS-induced mutations were recessive.

The screening was conducted in three batches, as tabulated in Table S1.

### Whole Genome Sequencing Mapping Analysis

The suppressor mutants were backcrossed with the wild-type N2 Bristol strain animals four times (96, 97). In the first three backcrosses, one F3 animal homozygous for *klp-7(tm2143)* mutation showing the suppression of ectopic extension phenotype similar to the original suppressor mutant was selected. In the fourth backcross, five to ten F3 animals were selected, one from each F2 plate to ensure the selection of unique recombinant animals in this step.

The progenies of the homozygous recombinant animals were pooled together to extract the genomic DNA by the isopropanol precipitation method (98). The library preparation and Illumina sequencing of genomic DNA was outsourced to the Genome Technology Access Centre at McDonnell Genome Institute (MGI), Washington University. They constructed KAPA PCR-free libraries for each sample and sequenced on 0.02 total of a NovaSeq S4 flowcell (300 XP) to obtain paired-end Fastq files.

We analyzed the whole genome sequencing data using the tools available on the online Galaxy platform (listed in Table S6) (99–101). The default settings were used for the analysis with each tool. The mapping by sequencing guide at https://mimodd.readthedocs.io/en/latest/nacreousmap.html (100, 102) was used to design the workflow for the analysis. The tools in Table S6 are listed in the order they were used for analysis.

### Sequencing of *citk-1(ok2328)* allele

For sequencing the mutant allele, we reverse transcribed the RNA isolated from the mutant animals. As previously described the about to be starved plates of mutant animals, which contained animals from all life stages, were washed with 1xM9 buffer thrice. The worm pellet was collected and stored at-80°C for RNA isolation. Using the Qiagen RNeasy Mini kit (no. 74104; Qiagen) RNA was isolated from the thawed pellet. The isolated RNA was treated with DNase I (Ambion’s DNA-free kit AM1906) to get rid of any genomic DNA contamination. Around 3-4 µg of this RNA was then reverse transcribed into cDNA using Superscript III Reverse Transcriptase (18080093). Wild-type cDNA was also isolated and processed as a control. The sequencing of the mutated region of this cDNA was outsourced to ATGC sequencing facility at Regional Centre for Biotechnology, Faridabad, India. The sequencing confirmed that *ok2328* deletion mutation is a frameshift variant as the 16bp insert (CCTATACTTACCTCAG) is incorporated in cDNA in the place of deleted segment, which introduces a premature stop codon after 635^th^ amino-acid.

### Immobilization of worms for imaging experiments

The fourth larval staged worms (L4) expressing fluorescent reporters were immobilized and mounted on 5% agarose pads using 10 mM Levamisole (Sigma-Aldrich; L3008) for widefield fluorescence imaging. For confocal scanning worms were mounted on 7.5% agarose pads. We immobilized the worms for EBP-2::GFP reporter live imaging using 0.1-μm polystyrene bead suspension (Polysciences; 00876-15) on 15% agarose pads. The agarose was dissolved in 1XM9 buffer as described in previously used protocols (25).

### Widefield fluorescence imaging of mechanosensory neurons for quantifying developmental defects

The ALM and PLM neurons were visualized using a fluorescent reporter expressed under a mechanosensory neuron-specific reporter *Pmec-7::*GFP *(muIs32)* using a Leica DM5000 B fluorescent microscope with a 40X air objective (NA 0.75). The L4 staged animals were mounted on 5% agarose beds and immobilized in 10mM Levamisole (Sigma-Aldrich; L3008) in 1XM9 buffer. The morphology of ALM neurons in *klp-7*(*0*) and suppressor mutants was qualitatively scored based on their microscopic appearance. The posterior neurite of the ALM neuron was considered a mild ectopic extension if it ended before the vulva and a strong ectopic extension if it crossed the vulva. This method was used to calculate the percentage of neurites showing the ectopic extension phenotype.

### Image acquisition and analysis of neurite length of mechanosensory neurons using a point-scanning confocal microscope

The mechanosensory neurons were visualized by expressing a fluorescent protein under a touch neuron-specific promoter either Pmec-7::GFP (*muIs32*) or Pmec-4::RFP (*tbIs222*). All the phenotyping or imaging experiments were performed with L4 stage animals using a Nikon confocal microscope (A1HD25) with a 60X oil objective (NA 1.4). The 1.5% of a 488nm laser was used to image GFP-tagged animals and 5% of a 561nm laser was used to image RFP-tagged animals along with a simultaneous differential interference contrast image.

To quantify the absolute and relative lengths of the anterior and posterior processes of PLM we used ImageJ software (Figure 2D, 4B, S4B-E, S5E). It was used to draw segmented traces along the length of the process in fluorescent channel ImageJ and then measure the length of the trace/process. The measured length of the anterior process of PLM was normalized by the length of the trace drawn between the anal opening and the vulva of the corresponding animal, measured from the differential interference contrast image. To normalize the length of the posterior process of PLM its value was divided by the length of a trace drawn from the anal opening to the tip of the tail.

### 3-D Protein Modeling

The protein structure predictions were modeled using the Alphafold2 (63, 103, 104) on the Galaxy platform (https://usegalaxy.eu/) (99). The molecular graphics and analysis were performed using Chimera (105) (Figure S1C-D and S2E-F).

### Molecular cloning and the generation of new transgene

pNBR77 *pCR8::citk-1*. The *citk-1* cDNA was amplified using the primers AGR733 (5’ CCGAATTCGCCCTTATGAACGAATCAATATATACAACGC 3’) and AGR734 (5’ GTCGAATTCGCCCTTTTAGTTTTTGGATCTTTTCAATGT 3’) from yk2046g11, and the pCR8 cloning vector was linearized using the primers AGR607 (5’ AAGGGCGAATTCGAC 3’) and AGR608 (5’ AAGGGCGAATTCGAC 3’). The two linearized fragments were then recombined by infusion (Takara, 638947).

pNBRGWY160 *Pmec-4::citk-1.* It was constructed by single site LR recombination between the destination vector *Pmec-4*::GWY and *pCR8::citk-1*(pNBR52) cDNA entry clone using an LR clonase enzyme (Invitrogen;11791–020). It was injected at 5ng/µl in the mutant animals.

*Pttx-3*::RFP or *Pmyo-2::mCherry* was used as a coinjection marker at concentrations of 60 ng/µl or 2.5ng/µl, respectively, to generate transgenic strains. The injection mixture was brought to a total concentration of 110–120 ng/μl by adding pBluescript (pBSK) plasmid to the injection mixture.

### Blast Analysis

The reciprocal blast analysis was conducted for the *citk-1* protein sequence using the online NCBI protein blast platform (106, 107). The algorithm parameters were set at their default settings. The taxon-specific blast results were conducted for reciprocal blast analysis (106–108) (Figure S2C).

### Foldseek Analysis

The Foldseek structural analysis was performed at default parameters in 3Di/AA mode using the online Foldseek search server (62). The taxonomic filter was used for the taxon-specific blast results (109).

### Protein Sequence Alignment

The M-coffee server (https://tcoffee.crg.eu/apps/tcoffee/do:mcoffee) was used to align the protein sequences obtained from UniProt server. (45, 110). SIAS was used to compare sequence similarity and sequence identity (http://imed.med.ucm.es/Tools/sias.html).

### RNAi Experiments

As the double mutant of *citk-1(ok2328)* and *citk-2(ok2885)* could not be generated a neuron-specific knockdown of *citk-2* in *citk-1* mutant was performed using a well-established RNAi by feeding method in worms (52–54, 111) (Figure 3-4 and Figure S3 and Figure S4). We used an RNAi-sensitive strain *lin-15b(n744)X; tbIs222(Pmec-4::mCherry); shrSi2[Pmec-3::sid-1, Cbunc-119(+)]II,* with a neuron-specific expression of a transmembrane dsRNA transporter *sid-1* under the touch neuron-specific promoter (*Pmec-3)* (54) and a gentle touch neuron-specific fluorescent reporter *Pmec-4::RFP (tbIs222)*. The worms were fed on HT115 *E. coli* bacteria expressing dsRNA for knockdown of gene of interest. As a positive control, the knockdown of *dhc-1* and *ama-1* was tested in this strain. The RNAi of *dhc-1*(dynein heavy chain-1) successfully produced dead embryos (112) and RNAi of *ama-1*(encodes the large subunit of RNA polymerase-II) resulted in L1 (larval stage-1) arrest phenotype (113) with a 100% penetrance. The negative control for RNAi was the HT115 bacterial strain with empty plasmid L4440. The positive control for neuronal RNAi was *unc-22* (regulates muscle contraction and relaxation) (114–116), and for gentle touch neuron-specific RNAi was *mec-4* (allows ligand-gated Na channel activity and is required for gentle touch sensation) (117–119), which successfully resulted in twitching (100% penetrance) and loss of gentle touch sensation phenotype (60% penetrance), respectively. The bacterial strains carrying the dsRNA used in this work are a part of Ahringer’s library (52), which was purchased from Source Bioscience (3318_Cel_RNAi_complete). The bacterial culture and feeding were performed as described in a recent paper from our lab (54). Each dsRNA bacterial strain was grown in 4.5 ml of LB nutrient broth with Carbenicillin (50mg/ml) and Tetracycline (12.5mg/ml) antibiotics to an OD of 0.8-0.9. Once this OD was achieved, bacterial culture was centrifuged and the pellet was resuspended in 1 ml of 1XM9 containing 50mg/ml Carbenicillin, 12.5mg/ml Tetracycline, and 1.5mM IPTG. A 250µl of this bacterial suspension was used for seeding NGM plates (containing 50mg/ml Carbenicillin, 12.5mg/ml Tetracycline, and 1.5mM IPTG), followed by a 36 hrs. incubation at 25 C. 5-6 L3s were transferred to these plates and their progenies were imaged.

### Live imaging and analysis of EBP-2::GFP dynamics

We used ZEN2 Blue software on a Zeiss Observer Z1 microscope equipped with a Yokogawa CSU-XA1 spinning disk confocal head and a Photometric Evolve electron-multiplying charge-coupled device camera for fast time-lapse image acquisition. We captured the images in a frame of 120 × 120 μm^2^ (512 × 512 pixels2) using a 100×/1.46 NA objective at a frame rate of 2.64 frames per second for 2 minutes’ duration. We used 8.75mW of a 488-nm excitation laser to achieve the best signal-to-noise ratio for EBP-2::GFP.

For the rescue experiments in *citk-1(ok2328); juIs338* both the mutant animals and the rescued animals were imaged on Nikon Ti2 widefield fluorescence microscope equipped with an Andor Zyla VSC-02284 Camera with a 560MHz readout rate. We captured the images at a 400ms exposure rate, a gain of 4 and 2×2 binning in a frame of 1280 × 1060 pixels using a Plan Apo VC 100× Oil/1.4 NA objective at a frame rate of 2.44 frames per second for 2 minutes’ duration.

To analyze the captured time-lapse images, we generated kymographs using the Plugins/ KymoReslice-Wide/Max Intensity Projection tool (https://github.com/UU-cellbiology/KymoResliceWide) (120, 121) (https://imagej.nih.gov/ij/). The kymographs were generated from two 30-μm ROIs one from the PLM anterior (from the 30u distal end to the cell body) and the other from the posterior process (from the cell body to the distal end at 30u). Kymographs are distance v/s time plots with distance along the X-axis representing axon length in micrometers and time in seconds along the Y-axis. The movement of an EBP-2::GFP comet along the axon length in a kymograph is observed as a diagonal line. We annotated the diagonal lines moving away from the cell body as “plus-end-out” microtubules (+), and the lines moving towards the cell body were annotated as “minus-end-out” microtubules (-). The fraction polarity was calculated as the relative number of plus-end-out or minus-end-out growing microtubules to the total number of microtubules. The growth length and growth duration were calculated from the slope of these tracks, by using the analysis tool of ImageJ.

## Statistical analysis

We plotted and analyzed data using GraphPad Prism software version 9.0.2 and 9.5.1. The details of methods and statistical tests used for comparisons are given in figure legends. The contingency plots represent the percentage of neurites showing characterized phenotypes. The bar plots represent mean and standard error of mean (SEM) for respective samples. The X^2^ test (Fisher’s exact test) was used to compare proportions. The normality of data was assessed using Shapiro-Wilk test. For normally distributed data. For Normally distributed data Bartlett’s test was performed to compare variance across samples followed by Brown Forsythe and Welch ANOVA test as samples showed significant difference between variances. For samples that did not pass the normality test, the comparison was performed using Mann Whitney test for two groups or Kruskal Wallis with a post-hoc Dunn’s multiple comparison test for multiple groups. In each Figure panel, P values are provided for intergroup comparisons. The respective sample size (n) and the total number of biological replicates (N) is also mentioned in the figure legends.

## Acknowledgments

We are grateful to Yuji Kohara for sharing cDNAs. We thank National BioResource Project (NBRP), Japan, and Caenorhabditis Genetics Center (CGC) for providing strains. NBRP is supported by Japanese government and CGC is supported by the NIH Office of Research Infrastructure Programs (P40 OD010440). We acknowledge the contributions of Pankajam Thyagarajan, Dharmendra Puri, Sanskriti Swamy, Kapil Raj, Sreyashi Chandra, Akanksha Goyal, and Debopriya Roy in initiating the EMS mutagenesis. We thank Sandhya Koushika for sharing strains. For all the genotyping and cloning related sequencing we thank the sequencing facility at Genomics Facility of the Advanced Technology Platform Centre (ATPC) which is managed by the Regional Centre for Biotechnology (RCB) and is funded by the Department of Biotechnology (Grant No. BT.MED-II/ATPC/BSC/01/2010). This work is supported by the NBRC core fund from the Department of Biotechnology, DBT/Wellcome Trust India Alliance Senior Fellowship (Grant # IA/S/22/1/506243) to Anindya Ghosh-Roy.

## Author Contributions

Sunanda Sharma, and Anindya Ghosh-Roy designed experiments. Sunanda Sharma and Keerthana Ponniah did the genetic and whole genome sequencing mapping of the new mutants isolated in this study. Sunanda Sharma has performed all the experiments in this paper involving Insilco analysis, molecular cloning, crosses, imaging and data analysis. Sunanda Sharma and Anindya Ghosh-Roy wrote the manuscript.

## References

1. Rasband MN. The axon initial segment and the maintenance of neuronal polarity. Nature Reviews Neuroscience2010. p. 552–62.

2. Puri D, Ponniah K, Biswas K, Basu A, Dey S, Lundquist EA, et al. Wnt signaling establishes the microtubule polarity in neurons through regulation of kinesin-13. Journal of Cell Biology. 2021;220(9).

3. Witte H, Neukirchen D, Bradke F. Microtubule stabilization specifies initial neuronal polarization. Journal of Cell Biology. 2008;180(3):619–32.

4. Homma N, Takei Y, Tanaka Y, Nakata T, Terada S, Kikkawa M, et al. Kinesin superfamily protein 2A (KIF2A) functions in suppression of collateral branch extension. Cell. 2003;114(2):229–39.

5. Homma N, Zhou R, Naseer MI, Chaudhary AG, Al-Qahtani MH, Hirokawa N. KIF2A regulates the development of dentate granule cells and postnatal hippocampal wiring. eLife. 2018;7:e30935.

6. Yogev S, Shen K. Establishing Neuronal Polarity with Environmental and Intrinsic Mechanisms. Neuron. 2017;96(3):638–50.

7. Desai A, Mitchison TJ. MICROTUBULE POLYMERIZATION DYNAMICS. 1997.

8. Mitchison T, Kirschner M. Dynamic instability of microtubule growth. Nature. 1984;312:237–42.

9. Nogales E. STRUCTURAL INSIGHTS INTO MICROTUBULE FUNCTION. 2001.

10. Akhmanova A, Steinmetz MO. Control of microtubule organization and dynamics: Two ends in the limelight. Nature Reviews Molecular Cell Biology. 2015;16(12):711–26.

11. Yau KW, Schätzle P, Tortosa E, Pagès S, Holtmaat A, Kapitein LC, et al. Dendrites <EM>In Vitro</EM> and <EM>In Vivo</EM> Contain Microtubules of Opposite Polarity and Axon Formation Correlates with Uniform Plus-End-Out Microtubule Orientation. The Journal of Neuroscience. 2016;36(4):1071.

12. Nguyen MM, McCracken CJ, Milner ES, Goetschius DJ, Weiner AT, Long MK, et al. Gamma-tubulin controls neuronal microtubule polarity independently of Golgi outposts. Mol Biol Cell. 2014;25(13):2039–50.

13. Wilkes OR, Moore AW. Distinct Microtubule Organizing Center Mechanisms Combine to Generate Neuron Polarity and Arbor Complexity. Front Cell Neurosci. 2020;14:594199.

14. Sakakibara A, Ando R, Sapir T, Tanaka T. Microtubule dynamics in neuronal morphogenesis. Open Biol. 2013;3(7):130061.

15. Pongrakhananon V, Saito H, Hiver S, Abe T, Shioi G, Meng W, et al. CAMSAP3 maintains neuronal polarity through regulation of microtubule stability. Proceedings of the National Academy of Sciences of the United States of America. 2018;115(39):9750–5.

16. Yau KW, van Beuningen SF, Cunha-Ferreira I, Cloin BM, van Battum EY, Will L, et al. Microtubule minus-end binding protein CAMSAP2 controls axon specification and dendrite development. Neuron. 2014;82(5):1058–73.

17. Harterink M, Vocking K, Pan X, Soriano Jerez EM, Slenders L, Freal A, et al. TRIM46 Organizes Microtubule Fasciculation in the Axon Initial Segment. J Neurosci. 2019;39(25):4864–73.

18. Curcio M, Bradke F. Microtubule Organization in the Axon: TRIM46 Determines the Orientation. Neuron. 2015;88(6):1072–4.

19. van Beuningen Sam FB, Will L, Harterink M, Chazeau A, van Battum Eljo Y, Frias Cátia P, et al. TRIM46 Controls Neuronal Polarity and Axon Specification by Driving the Formation of Parallel Microtubule Arrays. Neuron. 2015;88(6):1208–26.

20. Maniar TA, Kaplan M, Wang GJ, Shen K, Wei L, Shaw JE, et al. UNC-33 (CRMP) and ankyrin organize microtubules and localize kinesin to polarize axon-dendrite sorting. Nat Neurosci. 2011;15(1):48–56.

21. Jiang K, Hua S, Mohan R, Grigoriev I, Yau KW, Liu Q, et al. Microtubule Minus-End Stabilization by Polymerization-Driven CAMSAP Deposition. Developmental Cell. 2014;28(3):295–309.

22. Stepanova T, Slemmer J, Hoogenraad CC, Lansbergen G, Dortland B, De Zeeuw CI, et al. Visualization of Microtubule Growth in Cultured Neurons via the Use of EB3-GFP (End-Binding Protein 3-Green Fluorescent Protein) Several microtubule binding proteins, including CLIP-170 (cytoplasmic linker protein-170), CLIP-115, and EB1 (end-binding protein 1), have been shown to associate specifically with the ends of growing microtubules in non-neuronal cells, thereby regulating microtubule dynamics and the binding of microtubules to. 2003.

23. Leterrier C, Vacher H, Fache M-P, d’Ortoli SA, Castets F, Autillo-Touati A, et al. End-binding proteins EB3 and EB1 link microtubules to ankyrin G in the axon initial segment. Proceedings of the National Academy of Sciences. 2011;108(21):8826-31.

24. Partoens M, De Meulemeester AS, Giong HK, Pham DH, Lee JS, de Witte PA, et al. Modeling Neurodevelopmental Disorders and Epilepsy Caused by Loss of Function of kif2a in Zebrafish. eNeuro. 2021;8(5).

25. Ghosh-Roy A, Goncharov A, Jin Y, Chisholm AD. Kinesin-13 and Tubulin Posttranslational Modifications Regulate Microtubule Growth in Axon Regeneration. Developmental Cell. 2012;23(4):716–28.

26. Dey S, Kumar N, Balakrishnan S, Koushika SP, Ghosh-Roy A. KLP-7/Kinesin-13 orchestrates axon-dendrite checkpoints for polarized trafficking in neurons. Molecular Biology of the Cell. 2024;35(9):ar115.

27. Bounoutas A, O’Hagan R, Chalfie M. The Multipurpose 15-Protofilament Microtubules in C. elegans Have Specific Roles in Mechanosensation. Current Biology. 2009;19(16):1362–7.

28. Chalfie M, Thomson JN. Structural and functional diversity in the neuronal microtubules of Caenorhabditis elegans. J Cell Biol. 1982;93(1):15–23.

29. Lumpkin EA, Marshall KL, Nelson AM. The cell biology of touch. J Cell Biol. 2010;191(2):237–48.

30. Zheng C, Diaz-Cuadros M, Nguyen KCQ, Hall DH, Chalfie M. Distinct effects of tubulin isotype mutations on neurite growth in Caenorhabditis elegans. Molecular Biology of the Cell. 2017;28(21):2786–801.

31. Lee HMT, Sayegh NY, Gayek AS, Jao SLJ, Chalfie M, Zheng C. Epistatic, synthetic, and balancing interactions among tubulin missense mutations affecting neurite growth in Caenorhabditis elegans. Mol Biol Cell. 2021;32(4):331–47.

32. Marcette JD, Chen JJ, Nonet ML. The Caenorhabditis elegans microtubule minus-end binding homolog PTRN-1 stabilizes synapses and neurites. Elife. 2014;3:e01637.

33. Richardson CE, Spilker KA, Cueva JG, Perrino J, Goodman MB, Shen K. PTRN-1, a microtubule minus end-binding CAMSAP homolog, promotes microtubule function in Caenorhabditis elegans neurons. Elife. 2014;3:e01498.

34. Neumann B, Hilliard MA. Loss of MEC-17 leads to microtubule instability and axonal degeneration. Cell Rep. 2014;6(1):93–103.

35. Lockhead D, Schwarz EM, O’Hagan R, Bellotti S, Krieg M, Barr MM, et al. The tubulin repertoire of C. elegans sensory neurons and its context-dependent role in process outgrowth. Mol Biol Cell. 2016;27(23):3717–28.

36. Cueva JG, Hsin J, Huang KC, Goodman MB. Posttranslational acetylation of alpha-tubulin constrains protofilament number in native microtubules. Curr Biol. 2012;22(12):1066–74.

37. Hilliard MA, Bargmann CI. Wnt signals and frizzled activity orient anterior-posterior axon outgrowth in C. elegans. Dev Cell. 2006;10(3):379–90.

38. Prasad BC, Clark SG. Wnt signaling establishes anteroposterior neuronal polarity and requires retromer in C. elegans. Development. 2006;133(9):1757–66.

39. Zheng C, Diaz-Cuadros M, Chalfie M. GEFs and Rac GTPases control directional specificity of neurite extension along the anterior–posterior axis. Proceedings of the National Academy of Sciences. 2016;113(25):6973–8.

40. Ackley BD. Wnt-signaling and planar cell polarity genes regulate axon guidance along the anteroposterior axis in C. elegans. Dev Neurobiol. 2014;74(8):781–96.

41. Boulin T, Hobert O. From genes to function: the C. elegans genetic toolbox. Wiley Interdiscip Rev Dev Biol. 2012;1(1):114–37.

42. Puri D, Sharma S, Samaddar S, Ravivarma S, Banerjee S, Ghosh-Roy A. Muscleblind-1 interacts with tubulin mRNAs to regulate the microtubule cytoskeleton in C. elegans mechanosensory neurons. PLoS Genetics. 2023;19(8).

43. Skop AR, Liu H, Yates J, Meyer BJ, Heald R. Dissection of the mammalian midbody proteome reveals conserved cytokinesis mechanisms. Science. 2004;305(5680):61-6.

44. Paysan-Lafosse T, Blum M, Chuguransky S, Grego T, Pinto BL, Salazar GA, et al. InterPro in 2022. Nucleic Acids Res. 2023;51(D1):D418-D27.

45. UniProt C. UniProt: the Universal Protein Knowledgebase in 2023. Nucleic Acids Res. 2023;51(D1):D523–D31.

46. Sayers EW, Bolton EE, Brister JR, Canese K, Chan J, Comeau DC, et al. Database resources of the national center for biotechnology information. Nucleic Acids Res. 2022;50(D1):D20–D6.

47. D’Avino PP. Citron kinase - renaissance of a neglected mitotic kinase. Journal of Cell Science: Company of Biologists Ltd; 2017. p. 1701–8.

48. Hsu JM, Chen CH, Chen YC, McDonald KL, Gurling M, Lee A, et al. Genetic analysis of a novel tubulin mutation that redirects synaptic vesicle targeting and causes neurite degeneration in C. elegans. PLoS Genet. 2014;10(11):e1004715.

49. Woods S, Coghlan A, Rivers D, Warnecke T, Jeffries SJ, Kwon T, et al. Duplication and retention biases of essential and non-essential genes revealed by systematic knockdown analyses. PLoS Genet. 2013;9(5):e1003330.

50. Timmons L, Fire A. Specific interference by ingested dsRNA. Nature. 1998;395(6705):854.

51. Fraser AG, Kamath RS, Zipperlen P, Martinez-Campos M, Sohrmann M, Ahringer J. Functional genomic analysis of C. elegans chromosome I by systematic RNA interference. Nature. 2000;408(6810):325-30.

52. Kamath RS, Martinez-Campos M, Zipperlen P, Fraser AG, Ahringer J. Effectiveness of specific RNA-mediated interference through ingested double-stranded RNA in Caenorhabditis elegans. Genome Biol. 2001;2(1):RESEARCH0002.

53. Kamath RS, Ahringer J. Genome-wide RNAi screening in Caenorhabditis elegans. Methods. 2003;30(4):313–21.

54. Singh P, Selvarasu K, Ghosh-Roy A. Optimization of RNAi efficiency in PVD neuron of C. elegans. PLoS One. 2024;19(3):e0298766.

55. Jiang K, Rezabkova L, Hua S, Liu Q, Capitani G, Altelaar AFM, et al. Microtubule minus-end regulation at spindle poles by an ASPM-katanin complex. Nature Cell Biology. 2017;19(5):480–92.

56. Gai M, Bianchi FT, Vagnoni C, Vernì F, Bonaccorsi S, Pasquero S, et al. ASPM and CITK regulate spindle orientation by affecting the dynamics of astral microtubules. EMBO reports. 2017;18(10):1870-.

57. Paramasivam M, Yoon JC, LoTurco JJ. ASPM and citron kinase co-localize to the midbody ring during cytokinesis. Cell Cycle. 2007;6(13):1605–12.

58. Szklarczyk D, Kirsch R, Koutrouli M, Nastou K, Mehryary F, Hachilif R, et al. The STRING database in 2023: protein-protein association networks and functional enrichment analyses for any sequenced genome of interest. Nucleic Acids Research. 2023;51(1 D):D638-D46.

59. Connolly AA, Osterberg V, Christensen S, Price M, Lu C, Chicas-Cruz K, et al. Caenorhabditis elegans oocyte meiotic spindle pole assembly requires microtubule severing and the calponin homology domain protein ASPM-1. Molecular Biology of the Cell. 2014;25(8):1298–311.

60. Piano F, Schetter AJ, Morton DG, Gunsalus KC, Reinke V, Kim SK, et al. Gene Clustering Based on RNAi Phenotypes of Ovary-Enriched Genes in C. elegans. Current Biology. 2002;12(22):1959–64.

61. Bianchi FT, Gai M, Berto GE, Di Cunto F. Of rings and spines: The multiple facets of Citron proteins in neural development. Small GTPases: Taylor and Francis Inc.; 2020. p. 122–30.

62. van Kempen M, Kim SS, Tumescheit C, Mirdita M, Lee J, Gilchrist CLM, et al. Fast and accurate protein structure search with Foldseek. Nature Biotechnology. 2023.

63. Yang Z, Zeng X, Zhao Y, Chen R. AlphaFold2 and its applications in the fields of biology and medicine. Signal Transduct Target Ther. 2023;8(1):115.

64. Ahmed Z, Douglas MR, Read ML, Berry M, Logan A. Citron kinase regulates axon growth through a pathway that converges on cofilin downstream of RhoA. Neurobiology of Disease. 2011;41(2):421–9.

65. Di Cunto F, Ferrara L, Curtetti R, Imarisio S, Guazzone S, Broccoli V, et al. Role of citron kinase in dendritic morphogenesis of cortical neurons. Brain Research Bulletin. 2003;60(4):319–27.

66. Camera P, Santos Da Silva J, Griffiths G, Giuffrida MG, Ferrara L, Schubert V, et al. Citron-N is a neuronal Rho-associated protein involved in Golgi organization through actin cytoskeleton regulation. Nature Cell Biology. 2003;5(12):1071–8.

67. Camera P, Schubert V, Pellegrino M, Berto G, Vercelli A, Muzzi P, et al. The RhoA-associated protein Citron-N controls dendritic spine maintenance by interacting with spine-associated Golgi compartments. EMBO Reports. 2008;9(4):384–92.

68. El-Amine N, Carim SC, Wernike D, Hickson GRX. Rho-dependent control of the Citron kinase, Sticky, drives midbody ring maturation. Mol Biol Cell. 2019;30(17):2185–204.

69. Gai M, Camera P, Dema A, Bianchi F, Berto G, Scarpa E, et al. Citron kinase controls abscission through RhoA and anillin. Mol Biol Cell. 2011;22(20):3768–78.

70. Bassi ZI, Verbrugghe KJ, Capalbo L, Gregory S, Montembault E, Glover DM, et al. Sticky/Citron kinase maintains proper RhoA localization at the cleavage site during cytokinesis. J Cell Biol. 2011;195(4):595–603.

71. Cunto FD, Imarisio S, Camera P, Boitani C, Altruda F, Silengo L. Essential role of citron kinase in cytokinesis of spermatogenic precursors. J Cell Sci. 2002;115(Pt 24):4819–26.

72. Kodba S, Chaigne A. Delayed abscission in animal cells – from development to defects. Journal of Cell Science. 2023;136(13):jcs260520.

73. Gruneberg U, Neef R, Li X, Chan EH, Chalamalasetty RB, Nigg EA, et al. KIF14 and citron kinase act together to promote efficient cytokinesis. J Cell Biol. 2006;172(3):363–72.

74. Pan H, Guan R, Zhao R, Ou G, Chen Z. Mechanistic insights into central spindle assembly mediated by the centralspindlin complex. Proc Natl Acad Sci U S A. 2021;118(40).

75. McKenzie C, Bassi ZI, Debski J, Gottardo M, Callaini G, Dadlez M, et al. Cross-regulation between Aurora B and Citron kinase controls midbody architecture in cytokinesis. Open Biol. 2016;6(3).

76. Dupraz S, Hilton BJ, Husch A, Santos TE, Coles CH, Stern S, et al. RhoA Controls Axon Extension Independent of Specification in the Developing Brain. Current Biology. 2019;29(22):3874–86.e9.

77. Omelchenko A, Firestein BL. Axonal Development: RhoA Restrains but Does Not Specify. Current Biology. 2019;29(22):R1179–R81.

78. Wang H, Tewari A, Einheber S, Salzer JL, Melendez-Vasquez CV. Myosin II has distinct functions in PNS and CNS myelin sheath formation. J Cell Biol. 2008;182(6):1171–84.

79. Rivera Alvarez J, Asselin L, Tilly P, Benoit R, Batisse C, Richert L, et al. The kinesin Kif21b regulates radial migration of cortical projection neurons through a non-canonical function on actin cytoskeleton. Cell Reports. 2023;42(7):112744.

80. Falnikar A, Baas PW. Neuronal migration re-purposes mechanisms of cytokinesis. Cell Cycle. 2013;12(23):3577–8.

81. Harding BN, Moccia A, Drunat S, Soukarieh O, Tubeuf H, Chitty LS, et al. Mutations in Citron Kinase Cause Recessive Microlissencephaly with Multinucleated Neurons. American Journal of Human Genetics. 2016;99(2):511–20.

82. Li H, Bielas SL, Zaki MS, Ismail S, Farfara D, Um K, et al. Biallelic Mutations in Citron Kinase Link Mitotic Cytokinesis to Human Primary Microcephaly. American Journal of Human Genetics. 2016;99(2):501–10.

83. Sarkisian MR, Li W, Cunto FD, D’Mello SR, Loturco JJ. Citron-Kinase, a Protein Essential to Cytokinesis in Neuronal Progenitors, Is Deleted in the Flathead Mutant Rat. 2002.

84. Razuvaeva AV, Graziadio L, Palumbo V, Pavlova GA, Popova JV, Pindyurin AV, et al. The Multiple Mitotic Roles of the ASPM Orthologous Proteins: Insight into the Etiology of ASPM-Dependent Microcephaly. Cells: MDPI; 2023.

85. Passemard S, Titomanlio L, Elmaleh M, Afenjar A, Alessandri JL, Andria G, et al. Expanding the clinical and neuroradiologic phenotype of primary microcephaly due to ASPM mutations. Neurology. 2009;73(12):962–9.

86. Tungadi EA, Ito A, Kiyomitsu T, Goshima G. Human microcephaly ASPM protein is a spindle pole-focusing factor that functions redundantly with CDK5RAP2. Journal of Cell Science. 2017;130(21):3676–84.

87. van der Voet M, Berends CWH, Perreault A, Nguyen-Ngoc T, Gönczy P, Vidal M, et al. NuMA-related LIN-5, ASPM-1, calmodulin and dynein promote meiotic spindle rotation independently of cortical LIN-5/GPR/Gα. Nature Cell Biology. 2009;11(3):269–77.

88. Buchman JJ, Durak O, Tsai LH. ASPM regulates Wnt signaling pathway activity in the developing brain. Genes Dev. 2011;25(18):1909–14.

89. Higgins J, Midgley C, Bergh AM, Bell SM, Askham JM, Roberts E, et al. Human ASPM participates in spindle organisation, spindle orientation and cytokinesis. BMC Cell Biology. 2010;11.

90. Holt MJE. The role of the autosomal recessive primary microcephaly protein ASPM: A protein involved in mitosis. 2021.

91. Chuang M, Goncharov A, Wang S, Oegema K, Jin Y, Chisholm Andrew D. The Microtubule Minus-End-Binding Protein Patronin/PTRN-1 Is Required for Axon Regeneration in C. elegans. Cell Reports. 2014;9(3):874–83.

92. Alves Domingos H, Green M, Ouzounidis VR, Finlayson C, Prevo B, Cheerambathur DK. The kinetochore protein KNL-1 regulates the actin cytoskeleton to control dendrite branching. Journal of Cell Biology. 2024;224(2):e202311147.

93. Akhmanova A, Steinmetz MO. Microtubule minus-end regulation at a glance. Journal of Cell Science: Company of Biologists Ltd; 2019.

94. Brenner S. The genetics of Caenorhabditis elegans. Genetics. 1974;77(1):71–94.

95. Mello C, Fire A. Chapter 19 DNA Transformation. In: Epstein HF, Shakes DC, editors. Methods in Cell Biology. 48: Academic Press; 1995. p. 451-82.

96. Zuryn S, Jarriault S. Deep sequencing strategies for mapping and identifying mutations from genetic screens. Worm. 2013;2(3):e25081-e.

97. Zuryn S, Le Gras S, Jamet K, Jarriault S. A strategy for direct mapping and identification of mutations by whole-genome sequencing. Genetics. 2010;186(1):427–30.

98. Lissemore JL, Lackner LL, Fedoriw GD, De Stasio EA. Isolation of Caenorhabditis elegans genomic DNA and detection of deletions in the unc-93 gene using PCR. Biochem Mol Biol Educ. 2005;33(3):219–26.

99. Galaxy C. The Galaxy platform for accessible, reproducible and collaborative biomedical analyses: 2022 update. Nucleic Acids Res. 2022;50(W1):W345–W51.

100. Maier W. Mapping and molecular identification of phenotype-causing mutations (Galaxy Training Materials) 2018 [Available from: https://training.galaxyproject.org/training-material/topics/variant-analysis/tutorials/mapping-by-sequencing/tutorial.html.

101. Doitsidou M, Jarriault S, Poole RJ. Next-generation sequencing-based approaches for mutation mapping and identification in Caenorhabditis elegans. Genetics: Genetics; 2016. p. 451–74.

102. Maier W, Moos K, Seifert M, Baumeister R. MiModD - Mutation Identification in Model Organism Genomes: SourceForge.net; 2014 2014.

103. Jumper J, Evans R, Pritzel A, Green T, Figurnov M, Ronneberger O, et al. Highly accurate protein structure prediction with AlphaFold. Nature. 2021;596(7873):583-9.

104. Richard E, Michael ON, Alexander P, Natasha A, Andrew S, Tim G, et al. Protein complex prediction with AlphaFold-Multimer. bioRxiv. 2022:2021.10.04.463034.

105. Pettersen EF, Goddard TD, Huang CC, Couch GS, Greenblatt DM, Meng EC, et al. UCSF Chimera--a visualization system for exploratory research and analysis. J Comput Chem. 2004;25(13):1605–12.

106. Camacho C, Coulouris G, Avagyan V, Ma N, Papadopoulos J, Bealer K, et al. BLAST+: architecture and applications. BMC Bioinformatics. 2009;10:421.

107. Altschul SF, Madden TL, Schäffer AA, Zhang J, Zhang Z, Miller W, et al. Gapped BLAST and PSI-BLAST: a new generation of protein database search programs. Oxford University Press; 1997.

108. Tatusov RL, Koonin EV, Lipman DJ. A genomic perspective on protein families. Science. 1997;278(5338):631-7.

109. Monzon V, Paysan-Lafosse T, Wood V, Bateman A. Reciprocal best structure hits: using AlphaFold models to discover distant homologues. Bioinform Adv. 2022;2(1):vbac072.

110. Wallace IM, O’Sullivan O, Higgins DG, Notredame C. M-Coffee: combining multiple sequence alignment methods with T-Coffee. Nucleic Acids Res. 2006;34(6):1692–9.

111. Conte D, MacNei LT, Walhout AJM, Mello CC. RNA Interference in Caenorhabditis elegans. Current Protocols in Molecular Biology. 2015;2015:26.3.1-.3.30.

112. Gonczy P, Pichler S, Kirkham M, Hyman AA. Cytoplasmic dynein is required for distinct aspects of MTOC positioning, including centrosome separation, in the one cell stage Caenorhabditis elegans embryo. J Cell Biol. 1999;147(1):135–50.

113. Firnhaber C, Hammarlund M. Neuron-specific feeding RNAi in C. elegans and its use in a screen for essential genes required for GABA neuron function. PLoS Genet. 2013;9(11):e1003921.

114. Waterston RH, Thomson JN, Brenner S. Mutants with altered muscle structure of Caenorhabditis elegans. Dev Biol. 1980;77(2):271–302.

115. Greene DN, Garcia T, Sutton RB, Gernert KM, Benian GM, Oberhauser AF. Single-molecule force spectroscopy reveals a stepwise unfolding of Caenorhabditis elegans giant protein kinase domains. Biophys J. 2008;95(3):1360–70.

116. Matsunaga Y, Qadota H, Furukawa M, Choe HH, Benian GM. Twitchin kinase interacts with MAPKAP kinase 2 in Caenorhabditis elegans striated muscle. Mol Biol Cell. 2015;26(11):2096–111.

117. Suzuki H, Kerr R, Bianchi L, Frokjaer-Jensen C, Slone D, Xue J, et al. In vivo imaging of C. elegans mechanosensory neurons demonstrates a specific role for the MEC-4 channel in the process of gentle touch sensation. Neuron. 2003;39(6):1005–17.

118. Bounoutas A, Chalfie M. Touch sensitivity in Caenorhabditis elegans. Pflugers Archiv European Journal of Physiology 2007. p. 691–702.

119. Calixto A, Chelur D, Topalidou I, Chen X, Chalfie M. Enhanced neuronal RNAi in C. elegans using SID-1. Nat Methods. 2010;7(7):554–9.

120. Katrukha E. ekatrukha/KymoResliceWide: KymoResliceWide 0.5. 0.5 ed: Zenodo; 2020.

121. Schindelin J, Arganda-Carreras I, Frise E, Kaynig V, Longair M, Pietzsch T, et al. Fiji: an open-source platform for biological-image analysis. Nat Methods. 2012;9(7):676–82.

