## Supplementary Figures for "The cytokinesis associated proteins CITK and ASPM-1 regulate neuronal microtubule dynamics and polarity in *C. elegans*"

**List of Supplementary figures and Tables:**

**Figure S1:** Isolated variants in *mec-7* (β-tubulin) and *mec-12* (α-tubulin) that suppress *klp-7(0).*

**Figure S2:** Mapping and characterization of *shr11* variant.

**Figure S3:** Microtubules are more dynamic in the PLM anterior process of *citk* mutant animals.

**Figure S4:** *citk-1* single mutants display a short PLM posterior process.

**Figure S5:** ASPM-1 mutants exhibit increased microtubule dynamics and can suppress *klp-7(0)* ectopic extension phenotype.

**Figure S6:** The sequence alignment of worm *citk-1* and *citk-2* with mammalian CIT proteins*.*

**Table S1:** No. of F1s screened.

**Table S2**: Identified SNPs in suppressor mutants.

**Table S3**: Descriptive results of blast analysis of *citk-1.*

**Table S4**: Strains used in this study.

**Table S5**: Transgenes generated.

**Table S6**: Tools used for analysis of WGS data.

**Supplementary Figures**

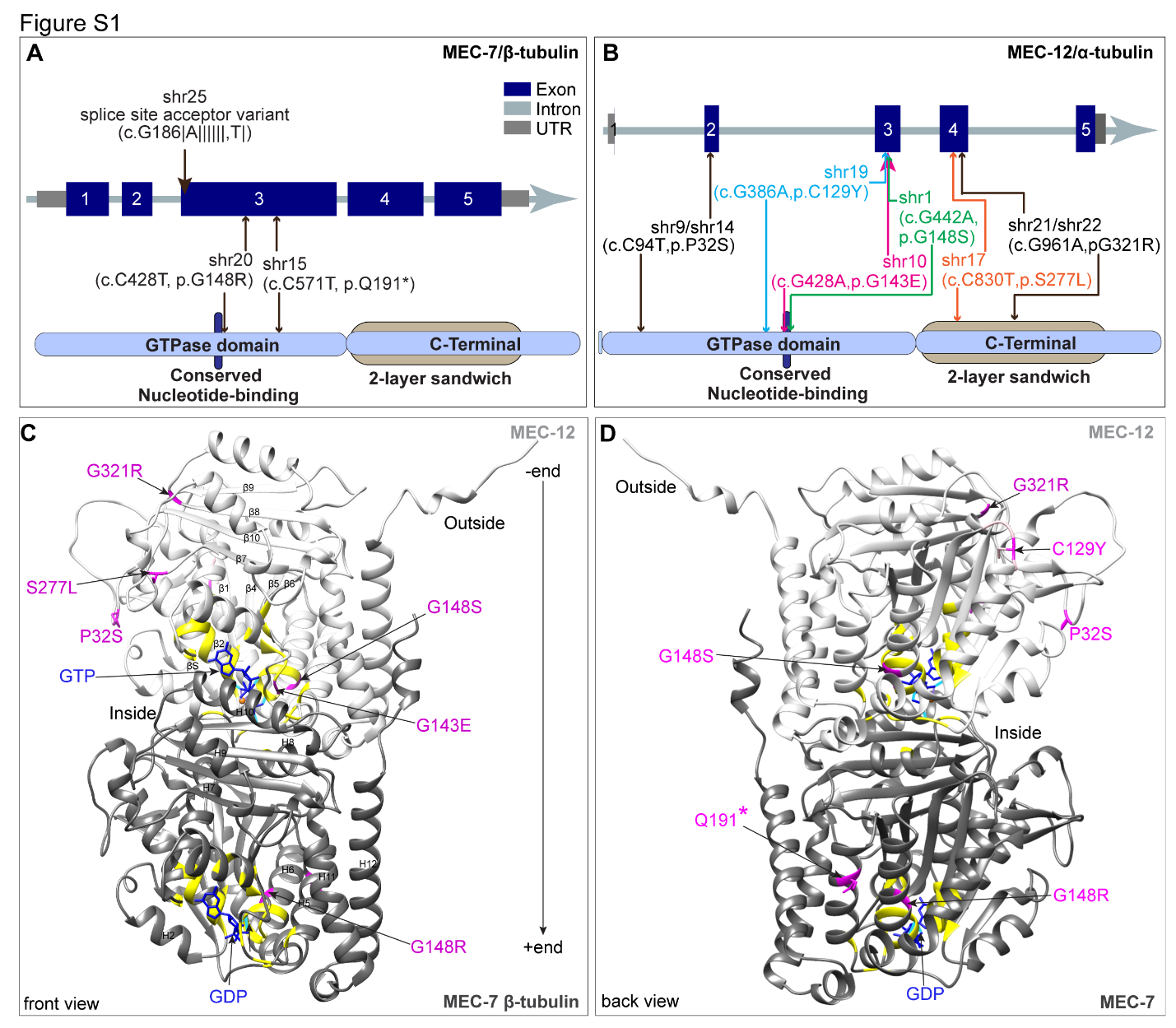

**Figure S1, related to Figure-1: Isolated variants in mec-7 (β-tubulin) and mec-12 (α-tubulin) that suppress klp-7(0).**

**(A-B)** Two-dimensional schematic illustration of the exon and intron of the *mec-7* and *mec-12* gene and protein structures with positions of identified candidate mutations labeled. The scale bar is in bp (base pairs). *mec-7* and *mec-12* gene. The *mec-7(shr20)* and *mec-7(shr15)* variants are in the nucleotide binding domain of the MEC-7 protein. The *shr25* is a splice site acceptor variant at the junction of intron-2 and exon-3 (A). The *mec-12(shr9), mec-12(shr19), mec-12(shr10)* and *mec-12(shr1)* variants are in the GTPase domain of protein and the *mec-12(shr17), and mec-12(shr21)* variants are in the C-terminal 2-layer sandwich domain (B). **(C-D)** Alphafold (1, 2) generated three-dimensional schematics of a MEC-7 and MEC-12 tubulin dimer with locations of *klp-7(0)* suppressor variants highlighted in magenta using Chimera (3) and rotated to display front and back views of the dimer. Several of these mutations are found in the nucleotide binding pocket of the tubulin. The *mec-7(shr15)* mutation introduces a premature stop codon. The *mec-12* mutation P32S is in the P loop and the G321R, S277L are present on the inter-dimer surface. These substitutions likely impact microtubule polymerization, nucleotide binding or heterodimer interactions.

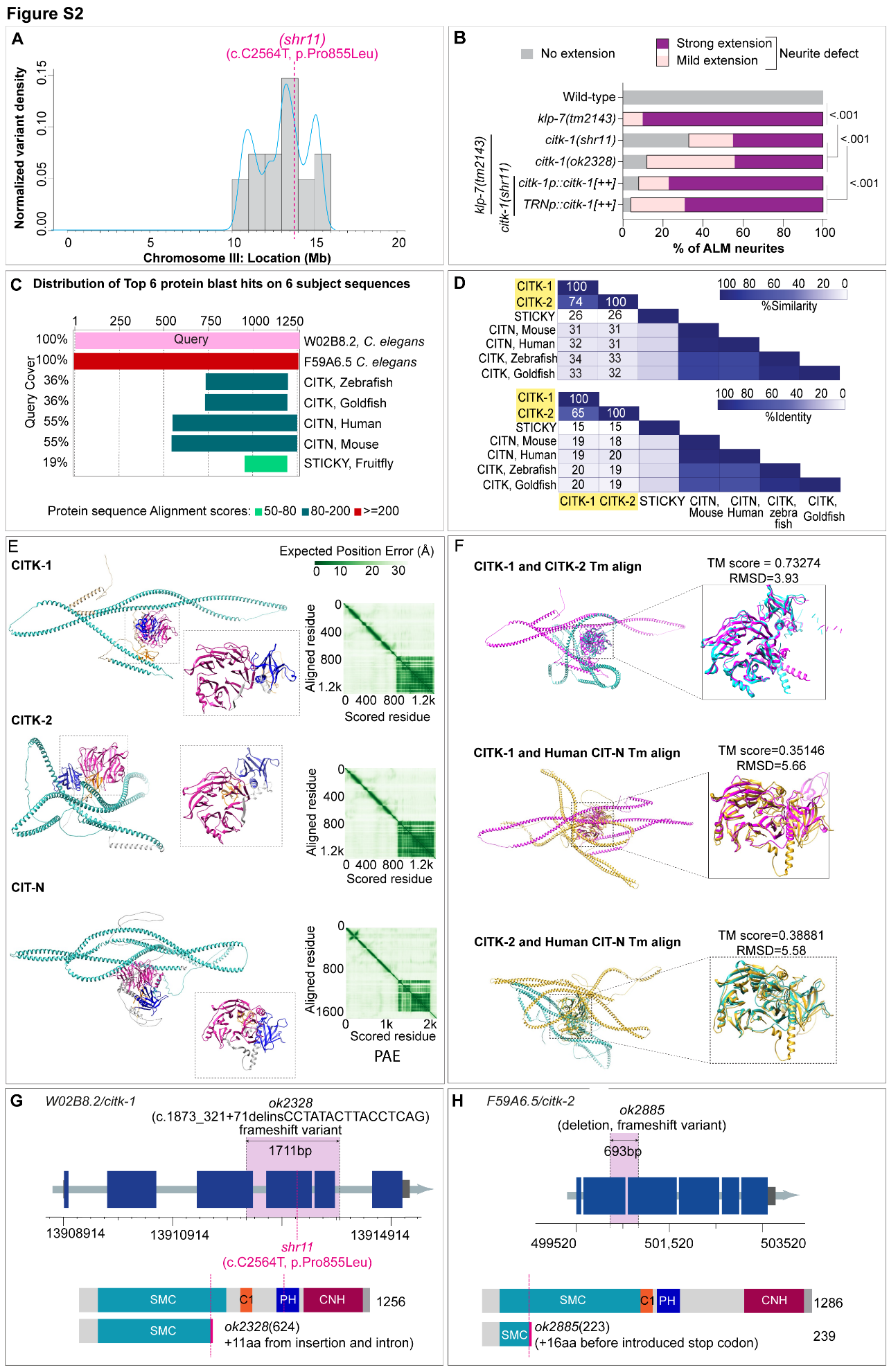

**Figure S2, related to Figure2: Mapping and characterization of *shr11* variant.**

**(A)** Normalized variant density plot of Chromosome II obtained by EMS density mapping analysis of *shr11* variant from the whole genome sequencing data using the method described in (4). The variant density is normalized as the number of EMS induced variants per 500Mb (gray bars) along the chromosomal length in Mb. It peaks at the genomic position linked to *shr11* variant. **(B)** Quantification of the percentage of ALM exhibiting ectopic extension phenotype in the wild-type, *klp-7(0),* *klp-7(0); citk-1(shr11),* *klp-7(0); citk-1(shr11); citk-1*(genomic DNA)[+] backgrounds, where *citk-1* (genomic DNA) is the fosmid WRM062dD08. N = 3–5 independent replicates, n (number of neurons) = 45-60. P values from 2x2 Fischer’s exact test. **(C)** Graphical overview of the distribution of top blast hits (5) on the query sequence, W02B8.2/CITK-1, represented by the pink bar. The hits are shown aligned to the regions of query, below in color coded bars. **(D)** Heat map showing the percentage similarity and percentage identity shared between citron rho interacting kinase proteins from different species. **(E)** The Alphafold predicted structures of CITK-1, CITK-2, and the kinase-less isoform of mammalian CIT (CIT-N). The inset shows the pleckstrin-homology domain (blue) and the citron-homology domain (magenta). The matrix on the right shows the expected position error in the predicted structures as given by Alphafold. **(F)** The Foldseek (6) predicted structural alignments and Tm alignment scores between CITK-1 and CITK-2, CITK-1 and CIT-N, and CITK-2 and CIT-N. Tm (Template modelling score) is a quantitative measure of protein structure similarity and lies between the values 0 and 1, where 1 indicates a perfect match. A Tm score below 0.3 indicates a random structural similarity (7). **(G-H)** The schematic of the exons and introns of the (G) citk-1 and (H) citk-2 gene and the genomic positions of (G) shr11, ok2328 and (H) ok2885 alleles in the gene locus. Below the gene structure is the schematic of (B) CITK 1 and (C) CITK-2 proteins and the deleted regions in the (G) ok2328 and (H) ok2885 alleles.

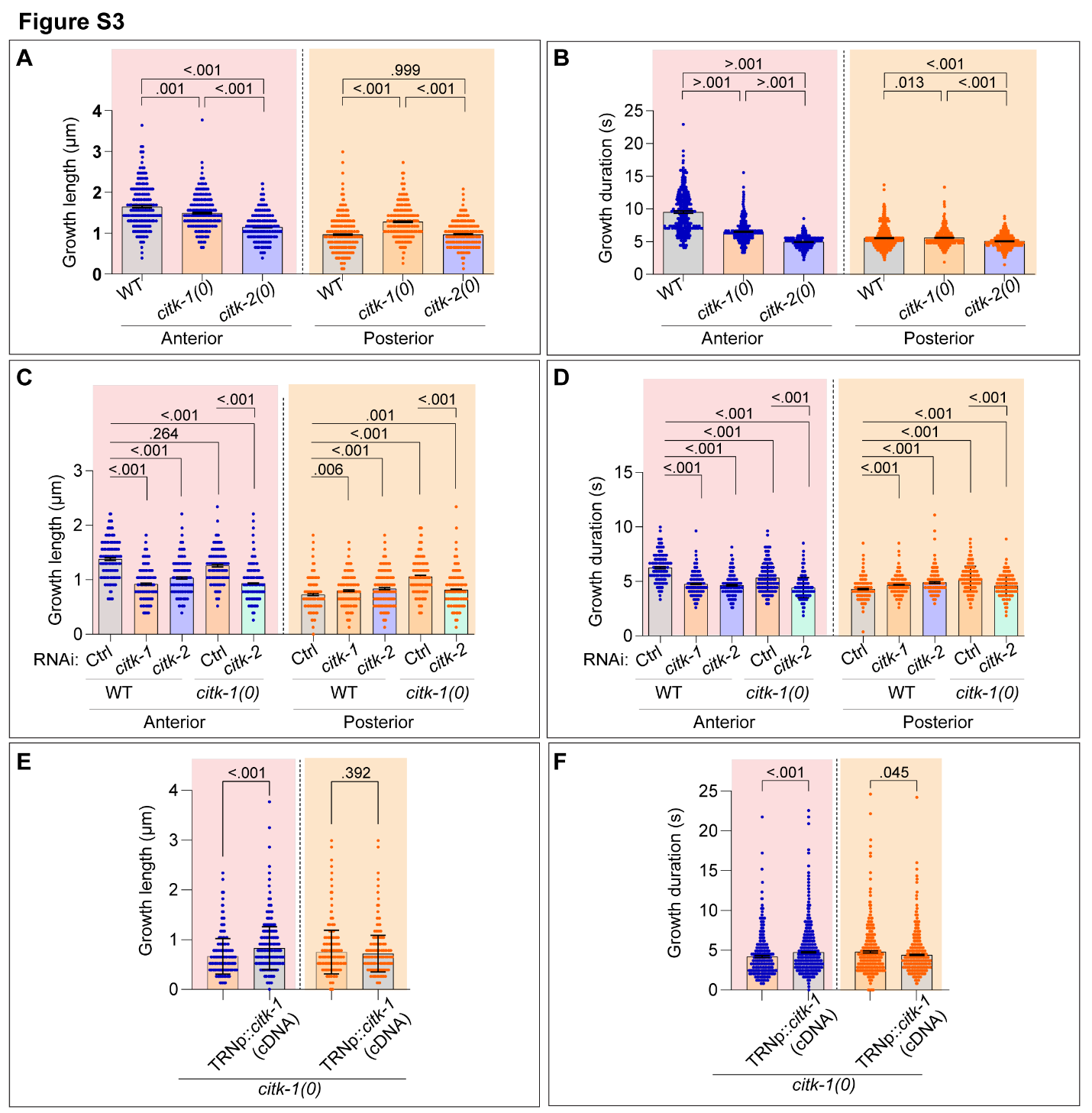

**Figure S3, related to figure 3: Microtubules are more dynamic in the PLM anterior process of *citk* mutant animals.**

Histograms depicting the growth length (A, C, E) and growth duration (B, D, F) of the EBP-2::GFP bound microtubules, quantified by determining the net pixel shift in the X and Y axis (of kymographs) respectively. (A-B) Quantifications from wild-type and citron kinase mutants. N = 3-4 independent replicates, n (number of tracks) = 343-676. (C-D) Quantifications from wild-type and *citk-1(0)* RNAi sensitive background animals fed for an RNAi mediated knockdown of *citk-1* or *citk-2* along with controls. N = 3 independent replicates, n (number of tracks) = 178-773. (E-F) Quantifications from *citk-1(0)* mutant animals and *citk-1(0)* rescue animals with a wild-type copy of *citk-1,* *Pmec-4::citk-1.* N = 3 independent replicates, n (number of tracks) = 506-1081. For A-D, Error bars represent SEM (Standard error mean), P values from Kruskal Wallis test followed by Dunn’s multiple comparison test, for E-F from Mann Whitney test. The groups separated by dotted lines were analyzed independent of each other.

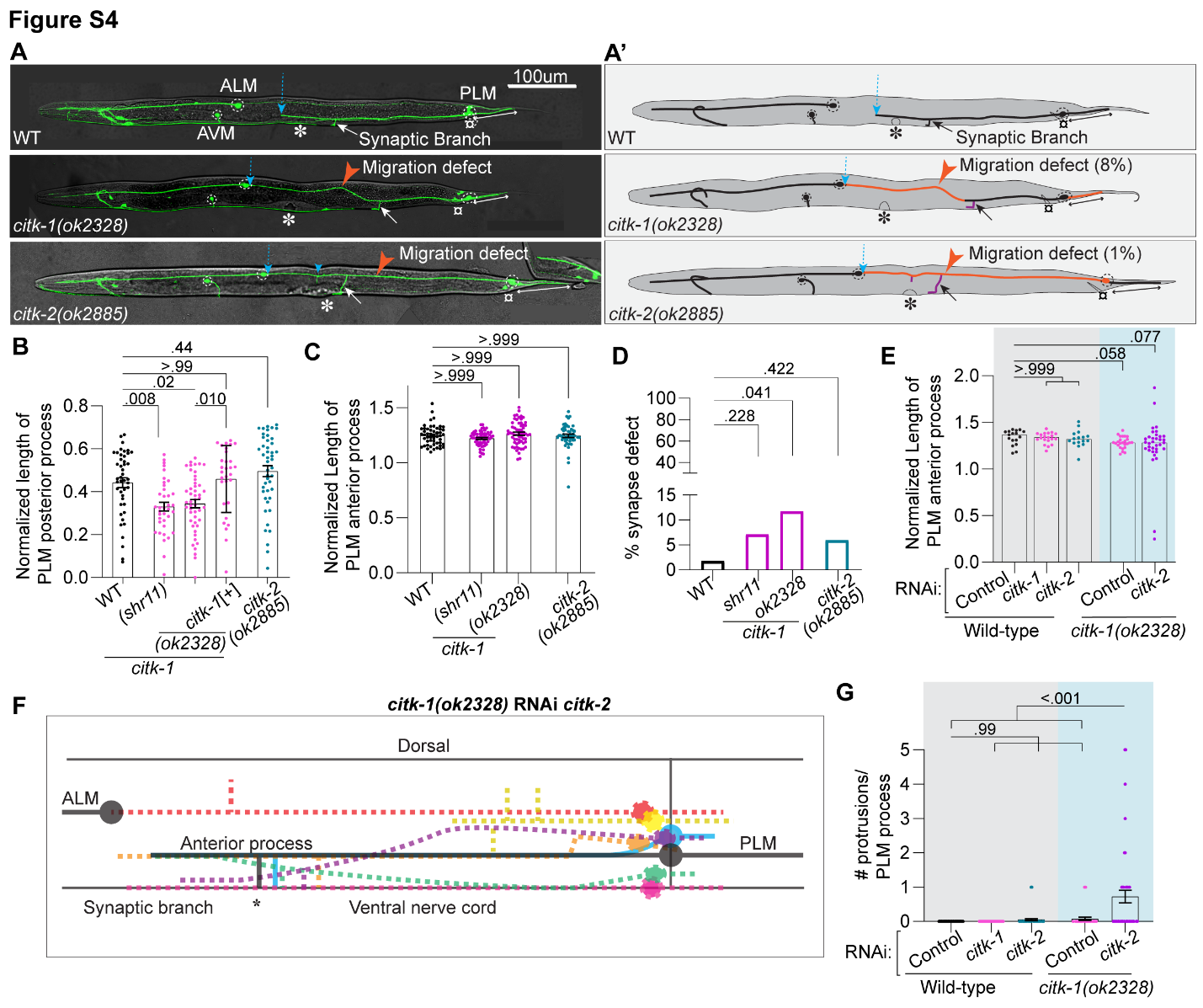

**Figure S4, Related to Figure-4: *citk-1* single mutants display a short PLM posterior process.**

**(A-A’)** Representative confocal images and schematics of the ALM and PLM mechanosensory neurons in L4 stage wild-type, *citk-1(ok2328)* and *citk-2(ok2885)* mutant animals showing cell positioning and migration defect. These neurons are labeled by *muIs32(Pmec-7::GFP)* reporter. PLM neuron grows laterally along the ventral edge of animal (positioned downwards). In 8% of *citk-1(0)* and merely 1% of *citk-2(0)* mutant animals the PLM neuron is misguided (pointed by a red arrowhead) towards the dorsal edge (positioned upwards) of the animal. A dotted blue arrow marks the distal tip of PLM anterior process. The white arrow points to the ventral synaptic branch. Blue arrowhead marks an ectopic protrusion. White double-headed arrow is drawn along the length of PLM posterior process. Vulva on the ventral side of animal is marked by an asterisk (*), and anus is marked by a generic symbol (¤). **(B-C)** Quantification of the normalized length of PLM posterior process (B) and anterior process (C) in the wild-type, *citk-1(shr11), citk-1(ok2328)* and *citk-2(ok2885)* animals. The PLM posterior process was also rescued by *shrEx514(Pcitk-1::citk-1),* extrachromosomal expression of a fosmid, WRM062dD08, containing complete genomic locus of *citk-1,* in *citk-1(ok2328)*mutant animals as quantified in (B). For B-D, N = 3-5 independent replicates, n (number of neurons) = 30-48. **(D)** Quantification of percentage of PLM neurons displaying a defect in the positioning or count of ventral synaptic branch in the wild-type, *citk-1(shr11), citk-1(ok2328)* and *citk-2(ok2885)* mutant animals. N = 3-5 independent replicates, n (number of neurons) = 51-68. (E) Quantification of the normalized length of PLM anterior process in the wild-type control and *citk-1(0)* RNAi sensitive background animals fed on bacteria expressing either empty vector (control) or *citk-1* or *citk-2* ds RNA for RNAi mediated knockdown. N = 3-5 independent replicates, n (number of neurons) = 17-34. (F) Schematic illustration of cell positioning and guidance defects observed in PLM neurons on knockdown of *citk-2* in *citk-1(0)* RNAi sensitive background as described in Figure 4E. The solid circles represent the positions of ALM and PLM cell bodies. The dorsal side of animal is towards the top of schematics and the anterior side is to the left. The vulval position is marked by an asterisk and the synaptic branch extends ventrally before vulva in the wild-type PLM (drawn in black solid line). The dotted lines represent different positions of PLM neurons as seen in *citk-1(0); citk-2*(RNAi kd). The neurons closer to the dorsal side above the WT PLM neuron are dorsally migrated and the neurons closer to the ventral nerve cord were called ventrally migrated. (G) Quantification of number of short ectopic protrusions/ PLM neuron anterior process in the wild-type control and *citk-1(0)* RNAi sensitive background animals fed on bacteria expressing either empty vector (control) or *citk-1* or *citk-2* ds RNA for RNAi mediated knockdown. N = 3-5 independent replicates, n (number of neurons) = 32-56. For B-C, E and G, Error bars represent SEM (Standard error mean), P values from Kruskal Wallis test followed by Dunn’s multiple comparison. For D, P values from 2x2 Fisher’s exact test.

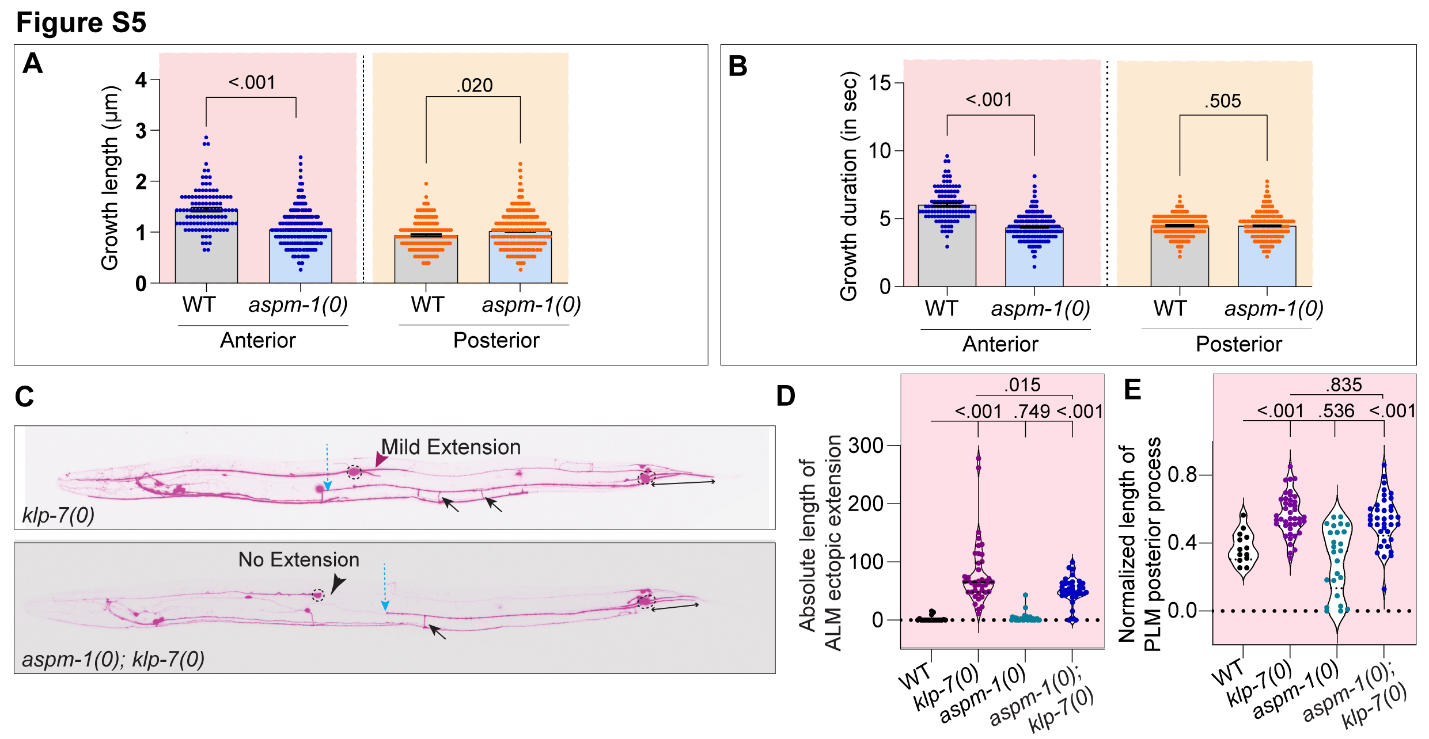

**Figure S5, Related to Figure 5: ASPM-1 mutants exhibit increased microtubule dynamics and can suppress *klp-7(0)* ectopic extension phenotype.**

**(A-B)** Histograms depicting the growth length (A) and growth duration (B) of the EBP-2::GFP bound microtubules in wild-type and *aspm-1(0)* mutants, quantified by determining the net pixel shift in the X and Y axis (of kymographs) respectively. The wild-type and *aspm-1(0)* animals were grown at non-permissive temperature of 25°C. N = 3-4 independent replicates, n (number of tracks) = 112-756. P values from Mann Whitney test. **(C)** Representative confocal images of ALM and PLM neurons in L4 staged *klp-7(0)* and *aspm-1(0); klp-7(0)* mutant animals grown at non-permissive temperature. The neurons are labeled by *muIs32(Pmec-7::GFP)* reporter. However, the image has been inverted in ImageJ for presentation purposes. The ALM extends two mild posterior extensions (pink arrowhead) in *klp-7(0)* animal. The *aspm-1(0); klp-7(0)* double mutant shows no extension. The blue dotted arrow marks the distal tip of anterior process of PLM neuron and the black double-headed arrow is drawn along the length of the PLM posterior process. Black solid arrows point to ventrally extended synaptic branches. **(E-F)** Quantification of the absolute and normalized lengths of ALM (E) and PLM (F) posterior processes, respectively in wild-type*,* *klp-7(0), aspm-1(0)* and *aspm-1(0); klp-7(0)* mutants. N = 3 and n (number of neurons) = 21-46. P values from Kruskal Wallis test followed by Dunn’s multiple comparison. Error bars represent SEM (Standard error mean).

**Figure S6**

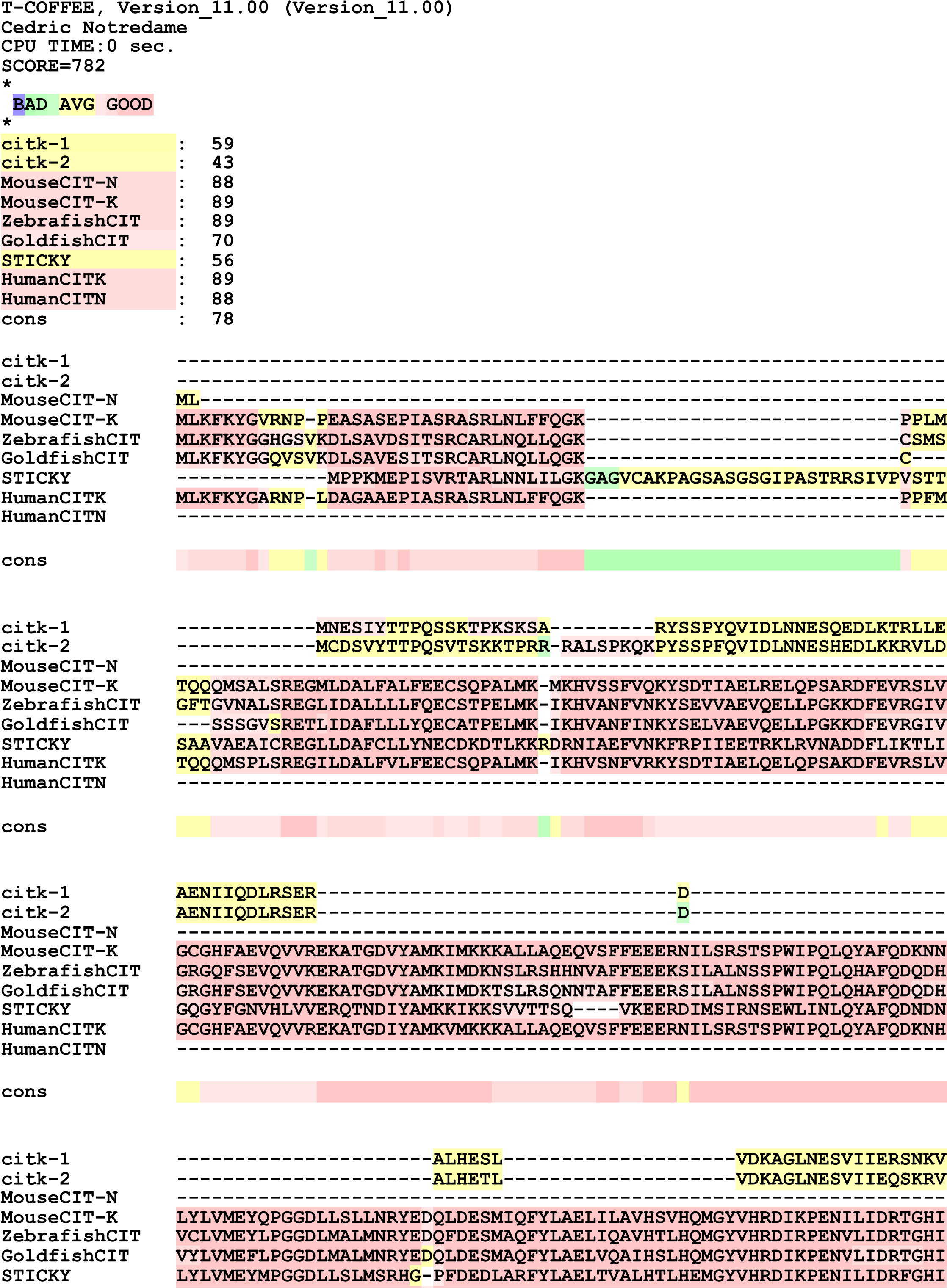

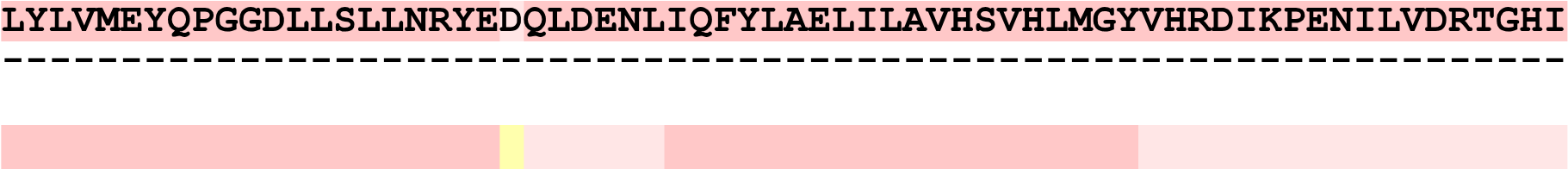

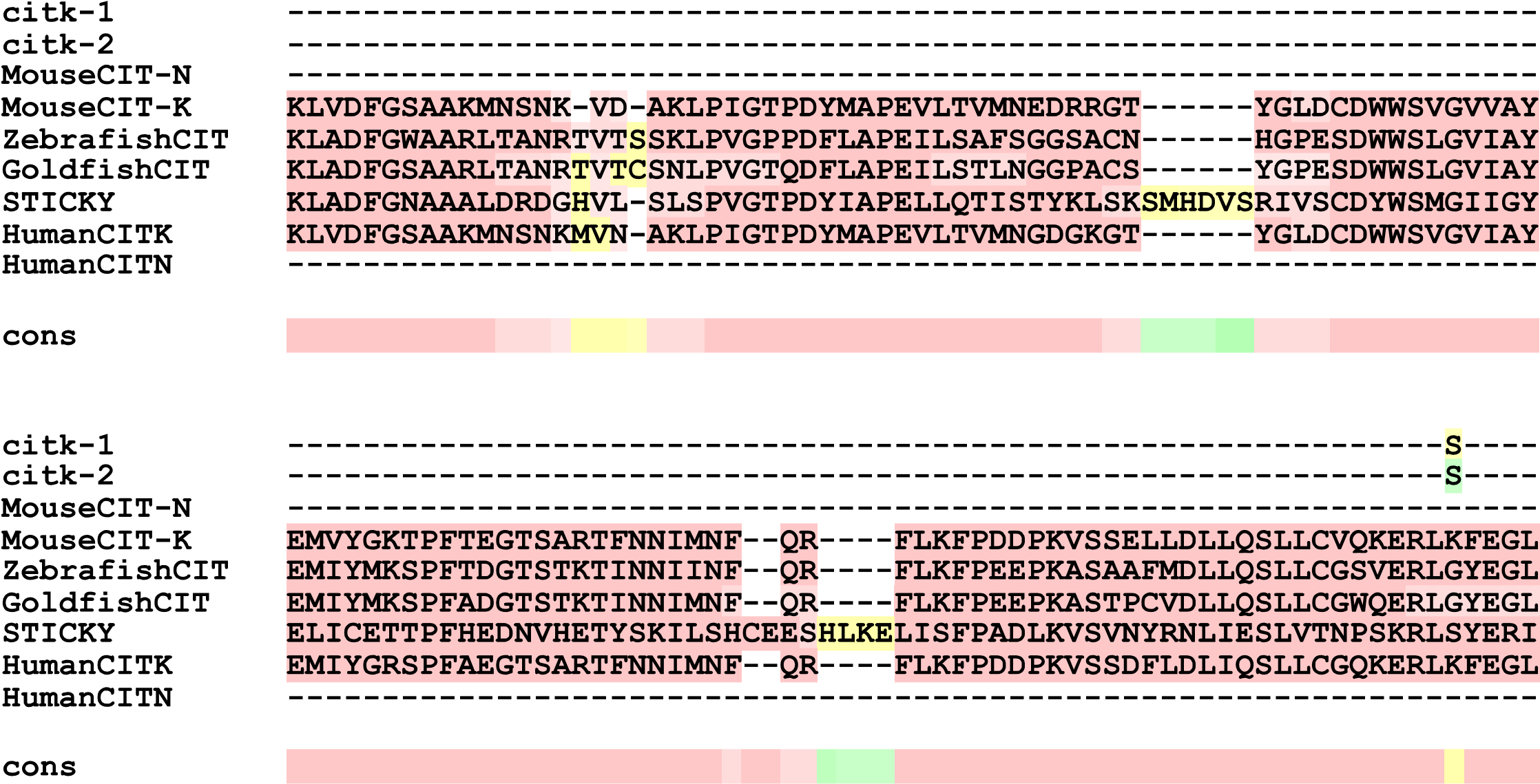

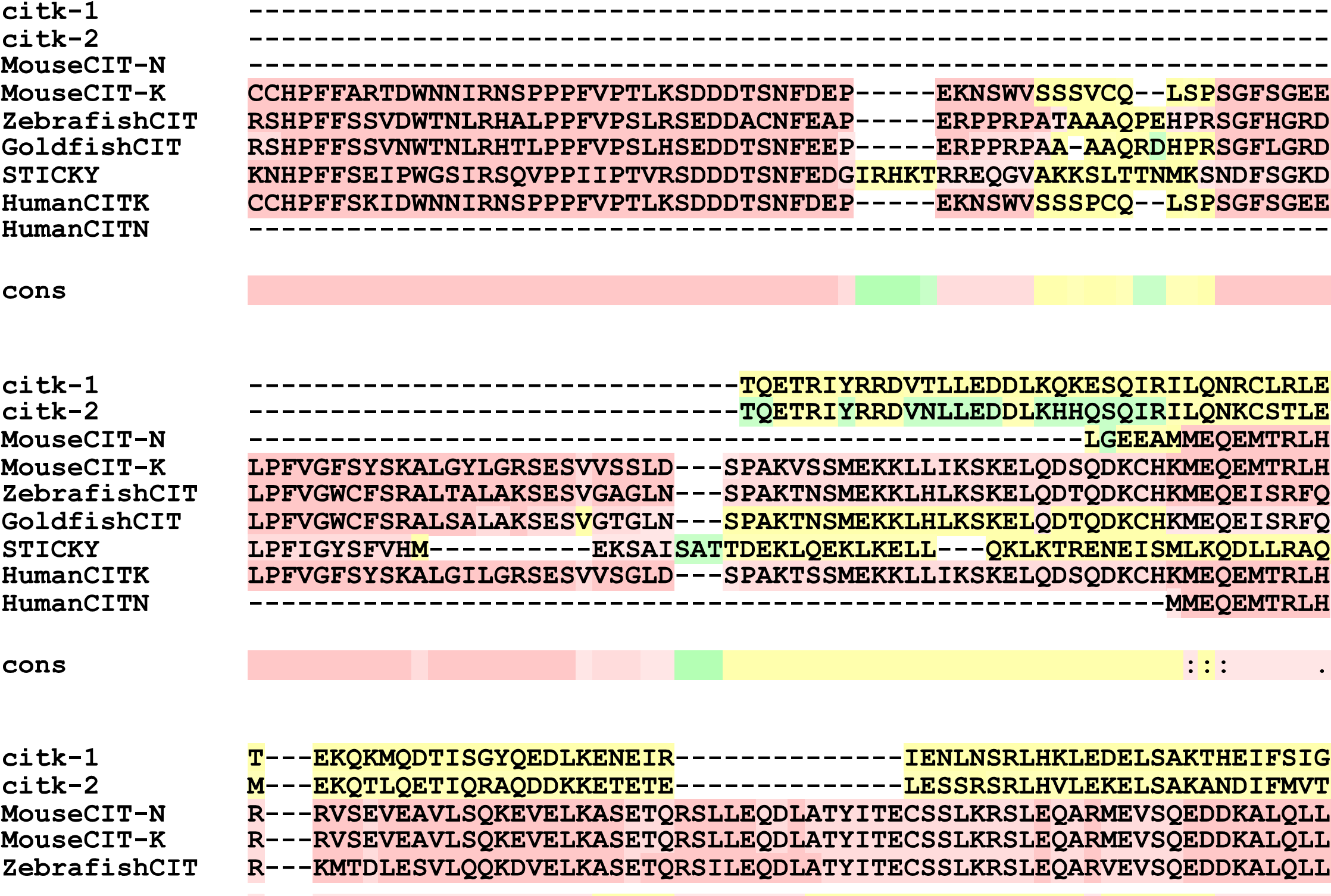

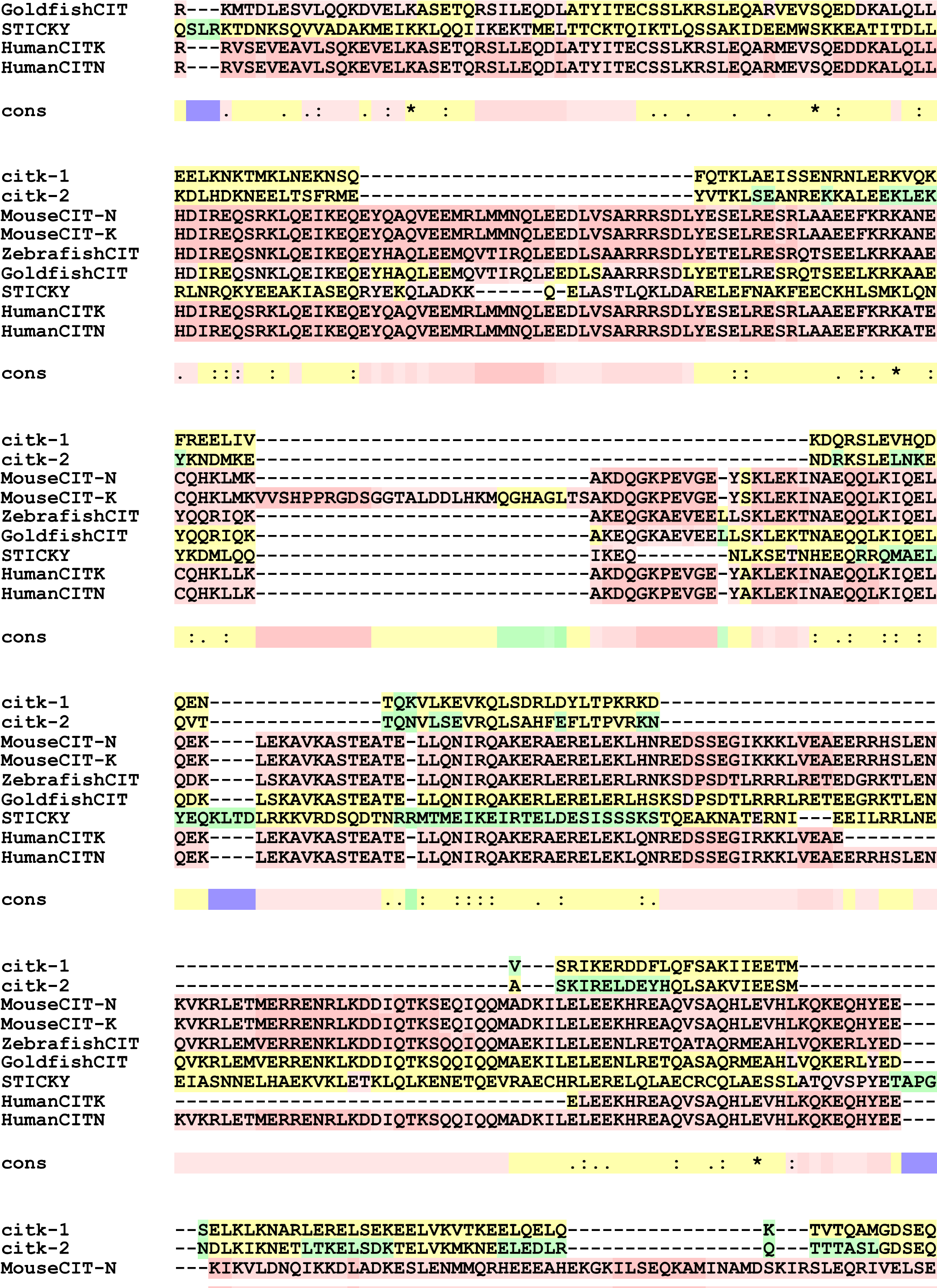

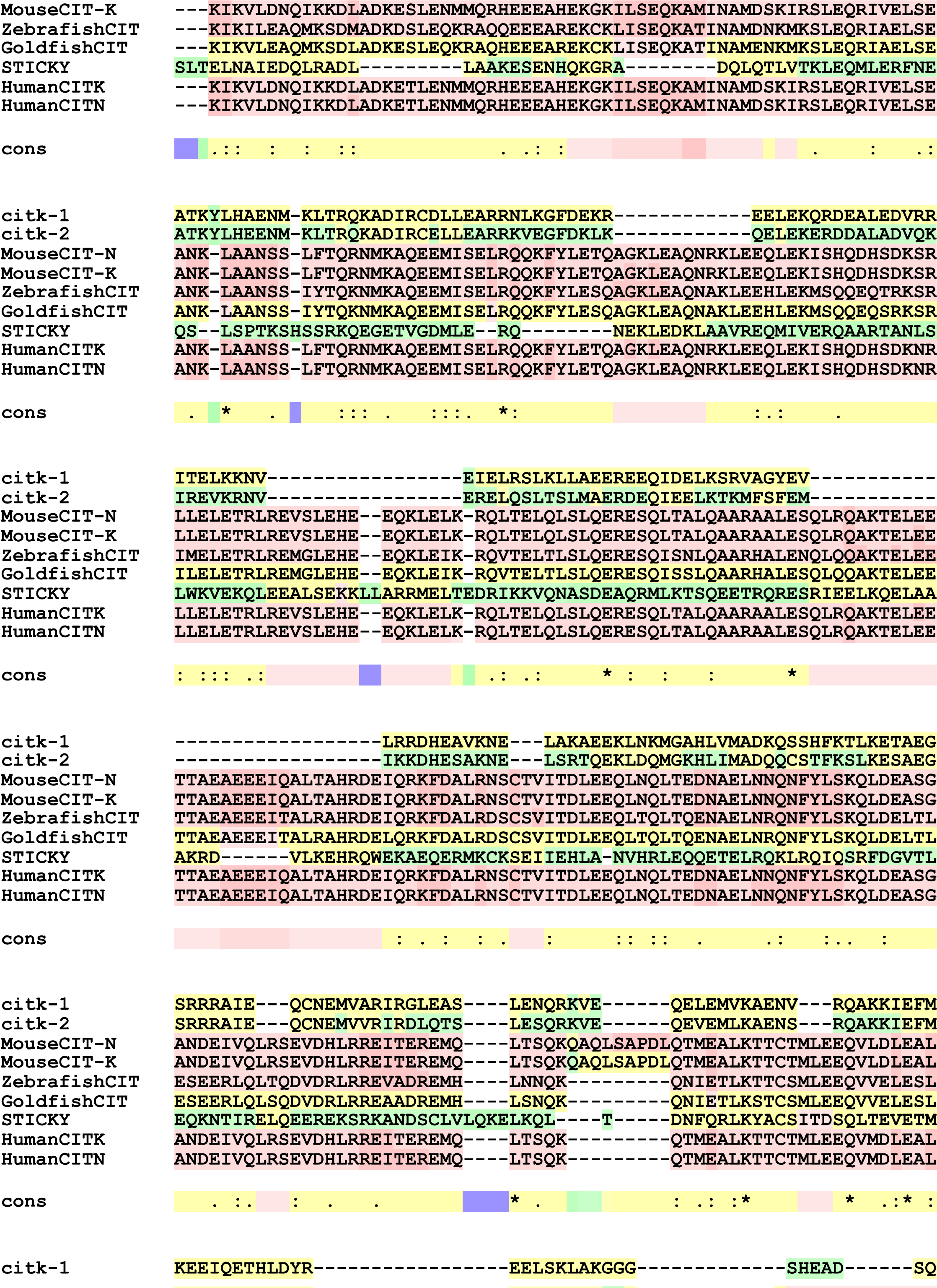

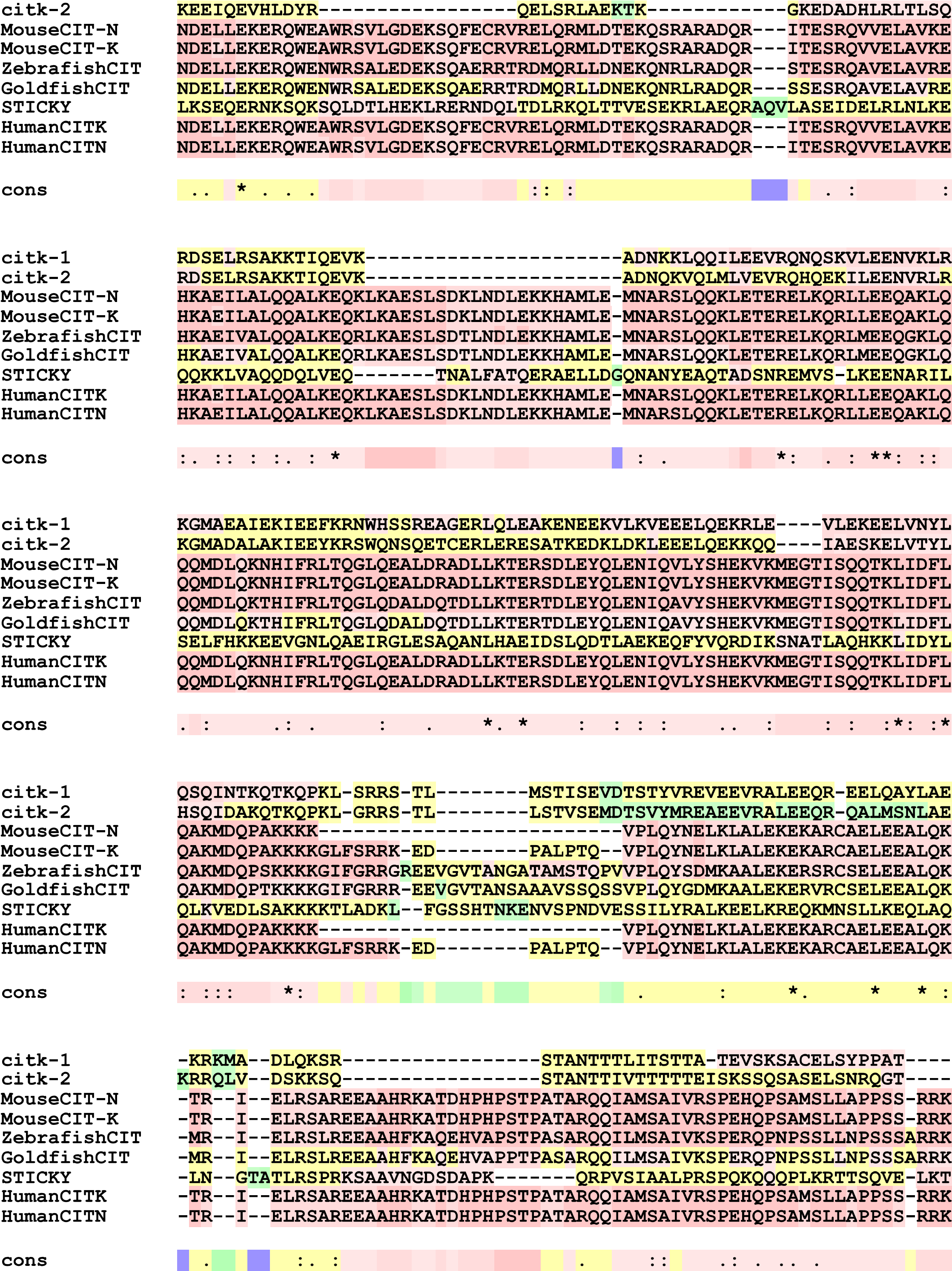

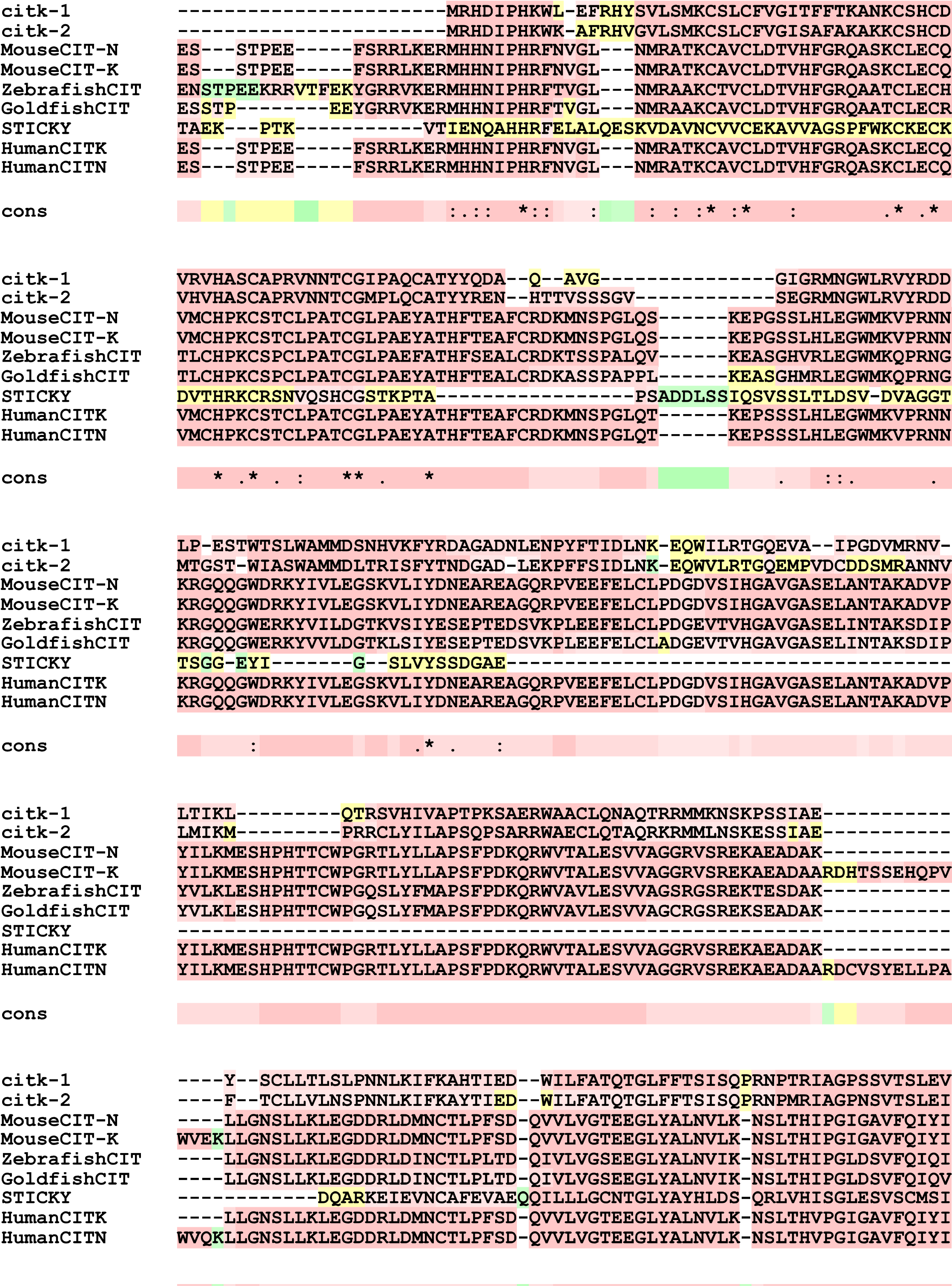

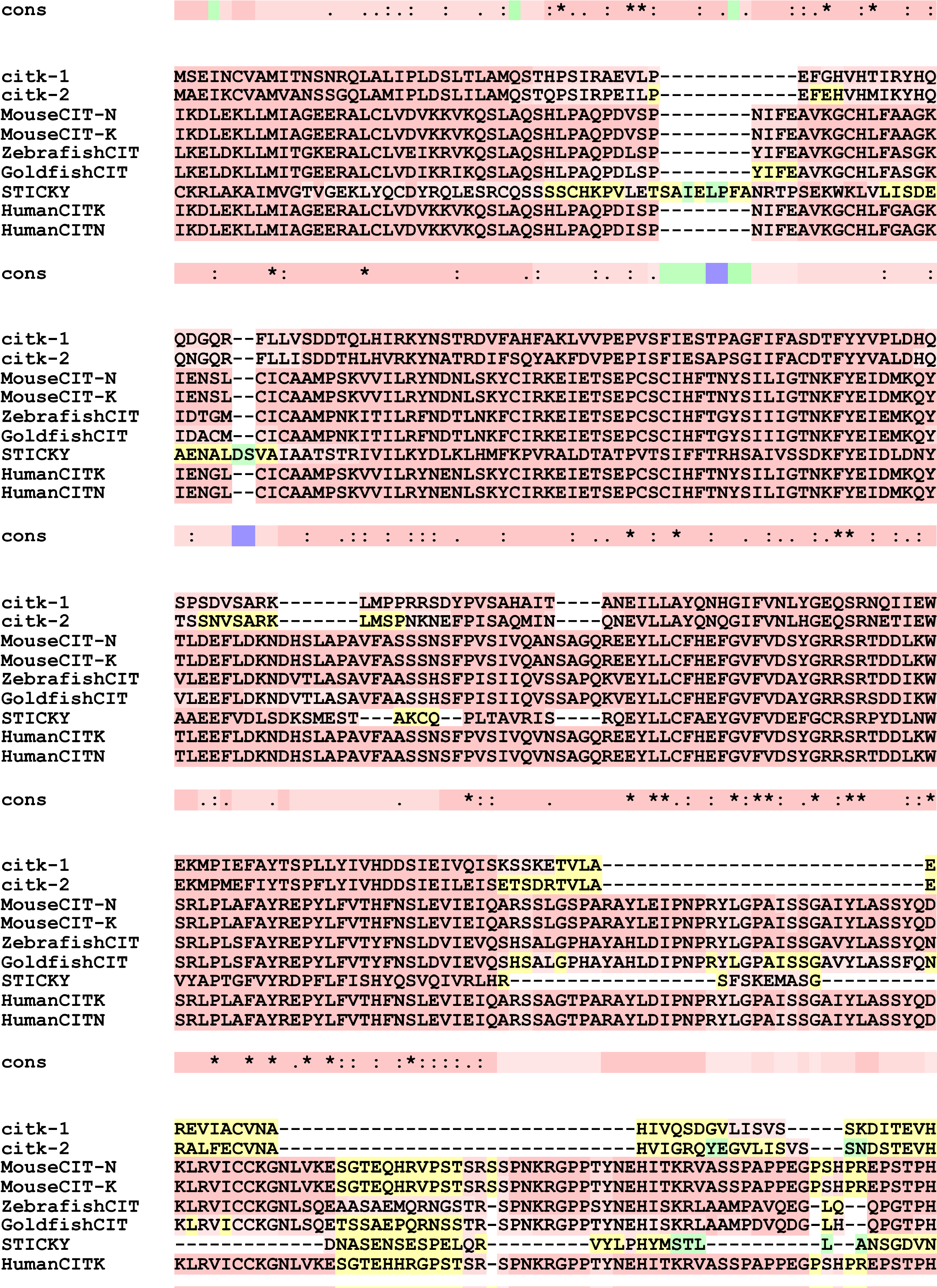

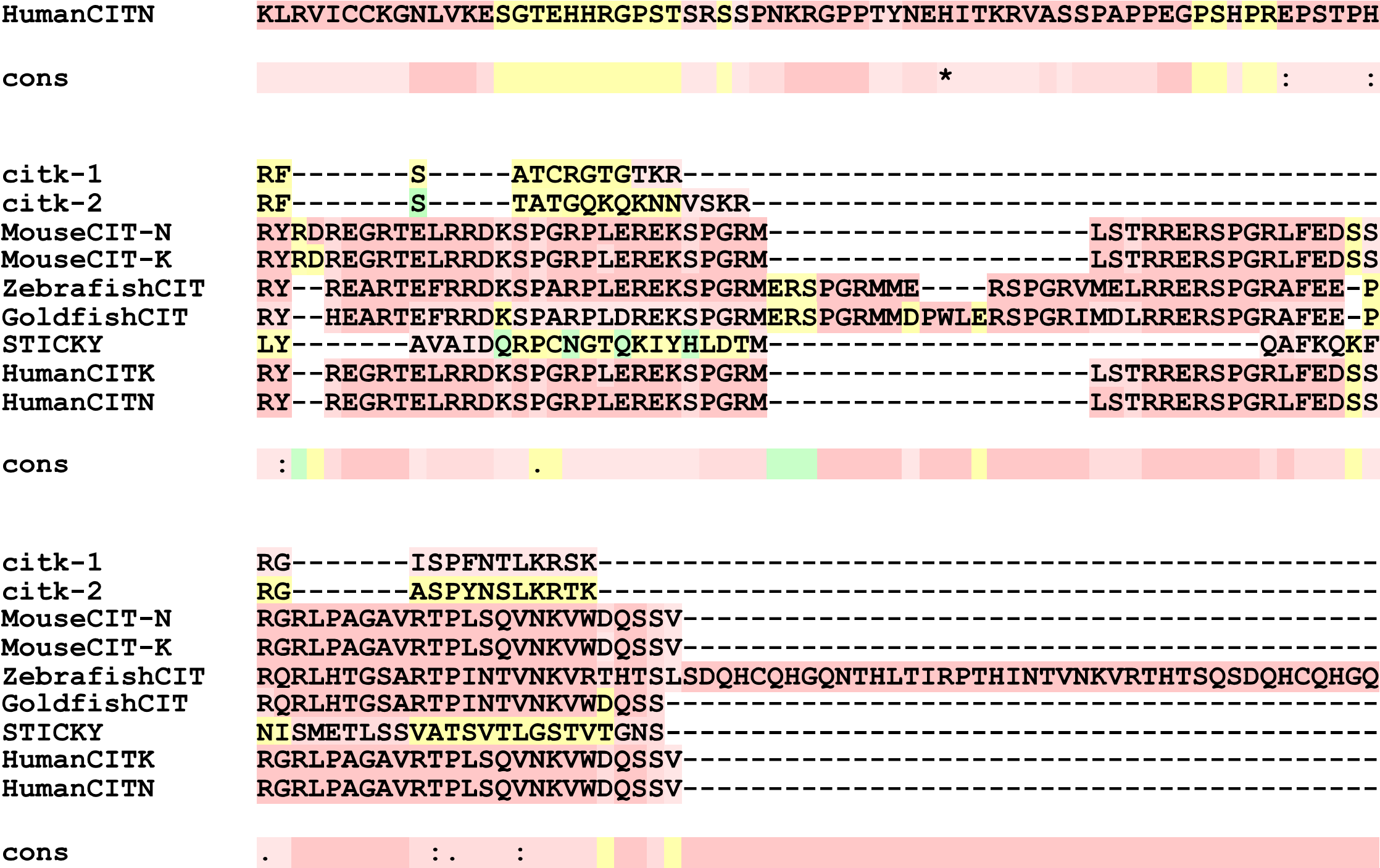

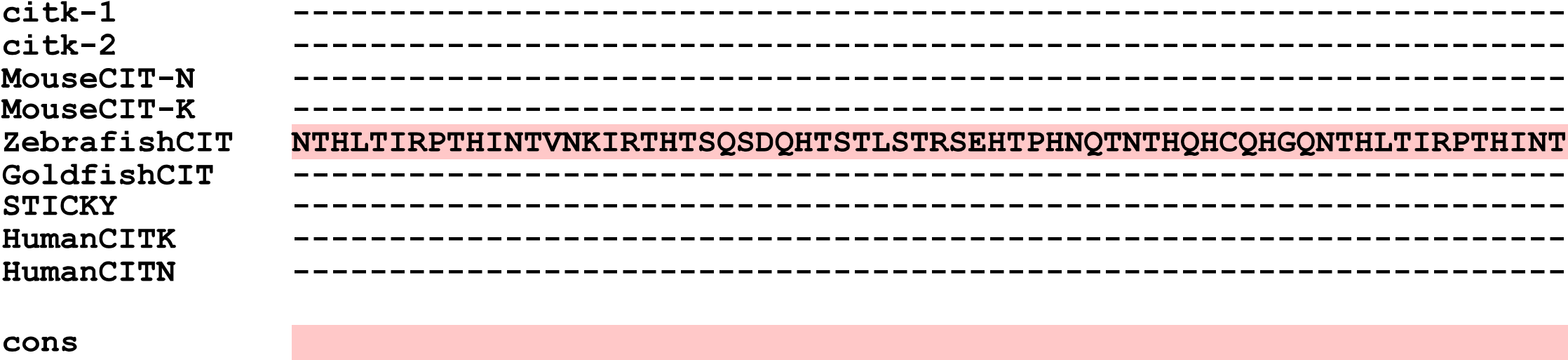

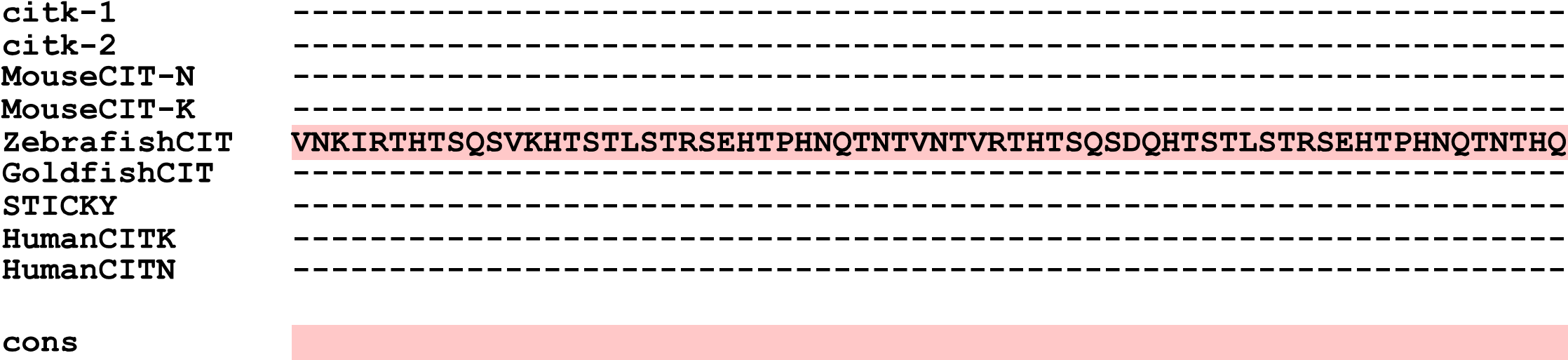

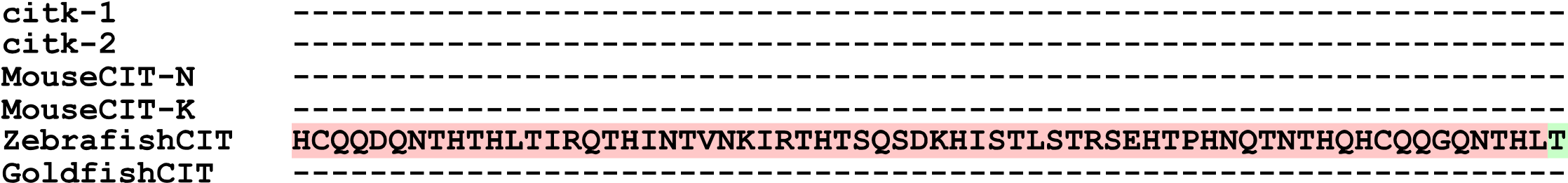

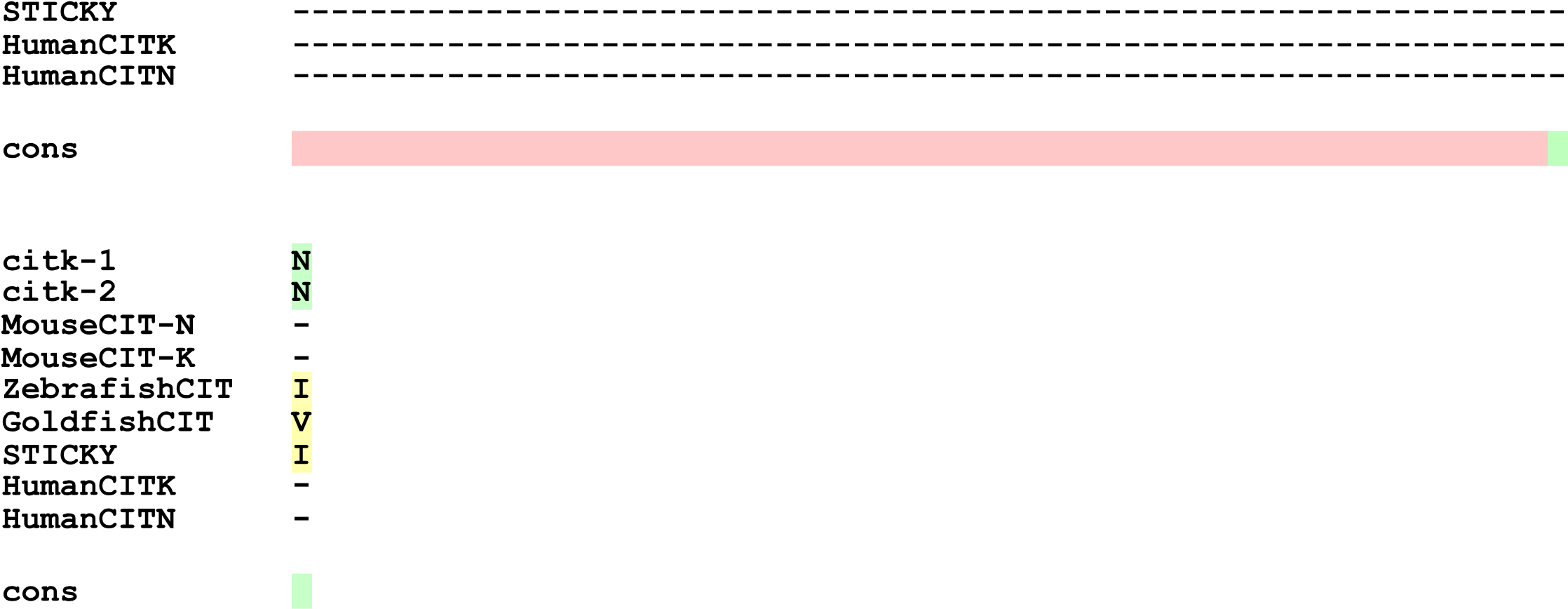

**Figure S6**: The sequence alignment of worm *citk-1* and *citk-2* with mammalian CIT proteins.

The sequence alignment of *citk-1/W02B8.2* and *citk-2/F59A6.5* proteins with the long (N-terminal kinase) and short isoforms (CITN-neuronal isoform) of Human and mouse CIT proteins and CIT orthologs of zebrafish, goldfish and Drosophila (sticky). The color scheme is used to denote conserved residues. Alignment was generated by T-coffee (https://tcoffee.crg.eu/apps/tcoffee/do:mcoffee).

References

1.	Jumper J, Evans R, Pritzel A, Green T, Figurnov M, Ronneberger O, et al. Highly accurate protein structure prediction with AlphaFold. Nature. 2021;596(7873):583-9.

2.	Richard E, Michael ON, Alexander P, Natasha A, Andrew S, Tim G, et al. Protein complex prediction with AlphaFold-Multimer. bioRxiv. 2022:2021.10.04.463034.

3.	Pettersen EF, Goddard TD, Huang CC, Couch GS, Greenblatt DM, Meng EC, et al. UCSF Chimera--a visualization system for exploratory research and analysis. J Comput Chem. 2004;25(13):1605-12.

4.	Doitsidou M, Jarriault S, Poole RJ. Next-generation sequencing-based approaches for mutation mapping and identification in Caenorhabditis elegans. Genetics: Genetics; 2016. p. 451-74.

5.	Altschul SF, Madden TL, Schäffer AA, Zhang J, Zhang Z, Miller W, et al. Gapped BLAST and PSI-BLAST: a new generation of protein database search programs. Oxford University Press; 1997.

6.	van Kempen M, Kim SS, Tumescheit C, Mirdita M, Lee J, Gilchrist CLM, et al. Fast and accurate protein structure search with Foldseek. Nature Biotechnology. 2023.

7.	Zhang Y, Skolnick J. TM-align: a protein structure alignment algorithm based on the TM-score. Nucleic Acids Res. 2005;33(7):2302-9.
